# Longitudinal immune dynamics of mild COVID-19 define signatures of recovery and persistence

**DOI:** 10.1101/2021.05.26.442666

**Authors:** Aarthi Talla, Suhas V. Vasaikar, Maria P. Lemos, Zoe Moodie, Mark-Phillip Lee Pebworth, Kathy E. Henderson, Kristen W. Cohen, Julie L. Czartoski, Lilin Lai, Mehul S. Suthar, Alexander T Heubeck, Palak C. Genge, Charles R. Roll, Morgan Weiss, Julian Reading, Nina Kondza, Hugh MacMillan, Olivia C. Fong, Zachary James Thomson, Lucas T. Graybuck, Lauren Y. Okada, Evan W. Newell, Ernest M. Coffey, Paul Meijer, Lynne A. Becker, Stephen C. De Rosa, Peter J. Skene, Troy R. Torgerson, Xiao-jun Li, Gregory Lee Szeto, M. Juliana McElrath, Thomas F. Bumol

## Abstract

SARS-CoV-2 has infected over 200 million and caused more than 4 million deaths to date. Most individuals (>80%) have mild symptoms and recover in the outpatient setting, but detailed studies of immune responses have focused primarily on moderate to severe COVID-19. We deeply profiled the longitudinal immune response in individuals with mild COVID-19 beginning with early time points post-infection (1-15 days) and proceeding through convalescence to >100 days after symptom onset. We correlated data from single cell analyses of peripheral blood cells, serum proteomics, virus-specific cellular and humoral immune responses, and clinical metadata. Acute infection was characterized by vigorous coordinated innate and adaptive immune activation that differed in character by age (young vs. old). We then characterized signals associated with recovery and convalescence to define and validate a new signature of inflammatory cytokines, gene expression, and chromatin accessibility that persists in individuals with post-acute sequelae of SARS-CoV-2 infection (PASC).

## Introduction

Severe acute respiratory syndrome-related coronavirus (SARS-CoV-2) is a novel, highly infectious human betacoronavirus that was first detected in late 2019, and has precipitated an ongoing pandemic that has infected >200 million people and killed over 4 million worldwide (Johns Hopkins COVID-19 dashboard, WHO dashboard). COVID-19, the clinical disease associated with SARS-CoV-2 infection, commonly presents with one or more respiratory symptoms, and may be accompanied by fatigue, fever, and often altered smell and taste; these can vary in severity with outcomes ranging from asymptomatic or mild to severe and fatal. There is substantial interindividual heterogeneity of severity, but more than 80% of infections are mild and individuals recover without hospitalization (Wu and McGoogan, 2020). Given the scale of worldwide infections, it is important to understand details of the normal, productive immune response to SARS-CoV-2 in mild COVID-19 infected individuals, who like those with more severe disease may experience persistent or recurrent symptoms. An estimated 30% to >70% of individuals with mild disease go on to develop post-acute sequelae of SARS-CoV2 infection (PASC or long COVID), an umbrella designation for clinical symptoms persisting weeks to months post-infection (Davis et al., 2020; Huang et al., 2021; Logue et al., 2021; Sudre et al., 2021).

The immune response to acute infection is critical to limit viral replication, activate innate immune cells, and efficiently prime virus-specific adaptive responses. Viral infections are rapidly sensed by pattern recognition receptors (PRRs) in infected cells and innate phagocytes, producing a key mediator of antiviral defense- interferons (IFNs). IFNs have delayed dynamics or are absent in severe COVID-19 (Galani et al., 2021; Hadjadj et al., 2020; Lucas et al., 2020) with multiple potential causes, including inefficient induction of type I and III IFNs in SARS-CoV-2-infected cells (Blanco-Melo et al., 2020), increased plasmacytoid dendritic cell (pDC) apoptosis and functional impairment (Arunachalam et al., 2020; Liu et al., 2021), and intrinsic type I IFN deficits such as somatic mutations or anti-IFN autoantibodies (Bastard et al., 2020; Wang et al., 2020a; Zhang et al., 2020). Other hallmarks of acute infection in severe COVID-19 include lymphopenia, impaired early IFN-stimulated genes (ISG) expression in monocytes, and robust plasmablast expansion and extrafollicular B cell responses (Mathew et al., 2020; Schulte-Schrepping et al., 2020; Woodruff et al., 2020). It is unclear if any of these features are shared with mild COVID-19, whether they are correlated with each other, and how they impact clinical outcome and convalescent immunity in this subset of disease. Understanding immune mechanisms of a successful acute infection response in mild COVID-19 and how these correlate with SARS-CoV-2-specific adaptive immune responses is critical to identifying biomarkers and potential therapeutic strategies to limit disease severity in SARS-CoV-2 infection.

Efficient resolution of inflammatory responses is necessary for recovery from acute infection. Failure to coordinate the kinetics or magnitude of inflammation can lead to dysregulated innate, cellular, and humoral immune responses (Carvalho et al., 2021; Schultze and Aschenbrenner, 2021; Sette and Crotty, 2021). A persistent inflammatory state is a characteristic of children and adults with multisystem inflammatory syndrome (MIS-C and MIS-A), including elevated inflammatory cytokines and chemokines in blood with similarities to secondary hemophagocytic lymphohistiocytosis (HLH) and cytokine release syndrome (CRS) (Consiglio et al., 2020; Fajgenbaum and June, 2020; Morris et al., 2020; Shaigany et al., 2020; Vella et al., 2020). In contrast, a typical mild COVID-19 trajectory and degree of interindividual heterogeneity remains poorly characterized. Detailed analysis of longitudinal trajectories is needed to identify key events for successful resolution of acute infection, and points of coordination between innate and adaptive immune responses over time. Analysis of immune response kinetics can also determine if persistence of symptoms in PASC is characterized by unique trajectories, and whether inflammatory responses are prolonged compared to successful convalescent cases.

To further our understanding of mild COVID-19, we performed longitudinal, deep immune phenotyping in SARS-CoV-2-infected participants and uninfected controls to define events coordinating innate and adaptive immunity, convalescent responses, and ongoing inflammation in the setting of persistent symptoms. Participants with a WHO ordinal severity score of 2 or 3 who did not require hospitalization were selected from a broader longitudinal COVID-19 cohort. Peripheral blood was sampled and analyzed longitudinally by 1) flow cytometry, 2) single cell RNAseq, 3) single cell ATACseq, 4) serum proteomics, 5) serology, and 6) *in vitro* anti-SARS-CoV-2-specific T and B cell responses. Samples were collected from at least 3 timepoints for every participant from 1-121 days post-symptom onset (PSO). We identified multimodal immune features over time that drive interindividual heterogeneity of SARS-CoV-2-specific immune responses, and correlated these with convalescent outcomes including levels of virus-specific memory B cells, antibodies, and occurrence of PASC. Serum protein profiling uncovered unique signatures that defined early acute infection and PASC. A validation cohort of individuals with PASC following a mild disease course confirmed and further defined these proteomic signatures. Integrative multi-omic analysis suggested key immune regulatory nodes in acute infection, the priming of adaptive immune responses, and PASC. These data provide a unique addition to existing and future COVID-19 related studies, and enable mechanistic hypothesis generation on mild disease progression and evolution to PASC. They highlight multiple targets for future validation in natural infection and vaccine-induced immunity as well as providing potential targets for immunomodulation in patients with persistent inflammatory disease after infection.

## Results

### Mild COVID-19 shows heterogeneity in clinical presentation and magnitude of acute and SARS-CoV-2-specific immune responses

We selected 20 SARS-CoV-2 PCR-positive participants with mild disease and 23 PCR-negative, uninfected controls from the Seattle metropolitan area. Of the COVID-19 participants, 18 had an initial sample collected less than 15 days from the onset of symptoms so these were selected for full longitudinal analysis (see **Methods** for inclusion criteria and quality control). The demographics and clinical features of this cohort are provided in **Supplementary Table 1**. Participants were categorized as younger (<40 years; n=14) and older (≥ 40 years; n=27) age groups (median age 29 vs. 57 years, respectively) (**Fig. 1A**). Each sample was processed in parallel into a multi-omic immunophenotyping pipeline including PBMCs analyzed by flow cytometry, scRNAseq, and scATACseq, while serum was analyzed by Olink proteomics (**Fig. 1B**). Each COVID-19 participant had 3-5 longitudinal study visits including, at a minimum: early acute infection (1-15 days PSO), late acute infection (16-30 days PSO), and post-acute COVID-19 (>30 days PSO) with a median follow-up of 81.5 days PSO (range: 33-121). Underlying comorbidities were documented and individuals with any symptom continuing or related to COVID-19 beyond 60 days were classified as PASC (**Fig. 1C**). Participants with post-acute illness symptoms that had a significant impact on activities of daily living were designated as sickest PASC. Uninfected controls had only a single study visit. Detailed symptom surveys were performed at each visit and cumulative results are summarized in **Fig. 1D**. Samples were also evaluated for SARS-CoV-2-specific adaptive immune responses by measuring IgG, IgM, and IgA antibody titers to spike (S) protein receptor binding domain (RBD), IgG to nucleocapsid (N) (Stamatatos et al., 2021), focus reduction neutralization assays against an infectious SARS-CoV-2 clone (Vanderheiden et al., 2020a), intracellular cytokine staining (ICS) to assess cytokine expression by activating CD4+ and CD8+ T cells with viral peptide pools covering SARS-CoV-2 structural proteins, and measurement of antigen-specific plasmablasts and memory B cells using S and RBD tetramers (**Fig. 1E-G**) (Cohen et al., 2021).

**Fig 1:**
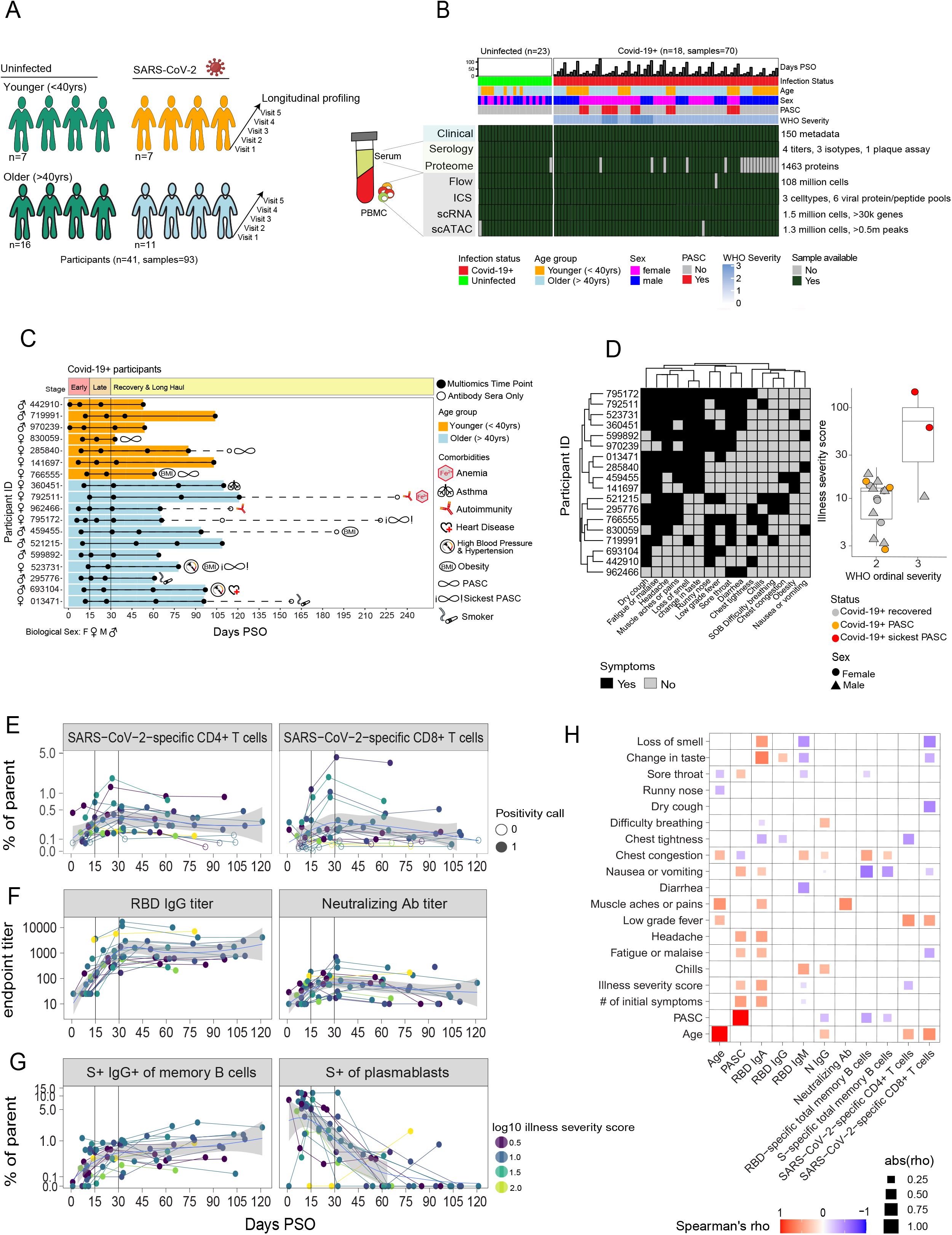
Overview of the longitudinal mild COVID-19 cohort from acute infection through convalescence. **A)** Cohort overview and participant demographics. Eighteen participants who had an initial blood draw within 15 days of developing symptoms (range 1-15 days) were followed longitudinally. **B)** Sample availability enumerated per assay type. PBMCs were analyzed by spectral flow cytometry, scRNAseq, scATACseq, and antigen-specific ICS assays. Serum was analyzed for SARS-CoV2 antibody serology or proteomics by Olink Explore 1536. Absent samples were due to limitations on material availability or timing. **C)** Longitudinal sampling timeline. PBMCs and serum were collected at 3-5 timepoints for each participant. Younger participants (top) are orange; older participants (bottom) are blue. Each Gantt is annotated with sex, documented comorbidities, and presentation of PASC. **D)** Self-reported symptoms occurring at any point during acute COVID-19 by participant. Illness severity score was calculated as described in Methods and shown vs. WHO ordinal severity. Each symbol represents one participant. Grey symbols indicate recovered COVID-19 participants, orange indicates PASC, and red indicates sickest PASC participants. **(E-G)** Cohort dynamics of SARS-CoV-2-specific adaptive immune responses: **E)** Aggregate frequencies of CD4+ and CD8+ T cells specific for E, M, N, S, and ORFs 3, 6a, 7a, 7b, and 8, **F)** titers of RBD IgG and neutralizing antibodies, and **G)** frequencies of S-specific of total plasmablasts and S-specific IgG+ of total memory B cells are shown over time for each participant with Loess smoothed curves overlaid and 95% confidence intervals shaded. Symbols for CD4+ and CD8+ T cell frequencies represent positive (filled) or negative (open) responses for each sample based on a positive response to any individual peptide pool stimulation determined using MIMOSA (Methods). **H)** Correlations between clinical metadata and estimates of SARS-CoV-2-specific immune responses. Day 30 estimates were used for all immune responses except neutralizing antibodies, which were estimated at day 42. Spearman’s rank coefficients were calculated for each pairwise combination, multiple comparisons corrected by Benjamini-Hochberg, and only correlations with adjusted p-values < 0.05 are displayed. The size of symbols represents the magnitude of correlations, while color represents magnitude and direction of positive (red) or negative (blue) correlation.

Illness severity was classified by participant reports of impact on activities of daily living for each day of acute illness (U S Department et al., 2017), Corrected Version 2.1). The WHO clinical progression scale was additionally used to classify each participant (Marshall et al., 2020). Despite being classified with a score of 2 or 3 on this WHO scale, mild COVID-19 participants had heterogeneous clinical presentations and disease courses. Sources of heterogeneity were primarily the spectrum of symptoms (**Fig. 1D**) and recovery time, ranging from 1-32 days of symptoms during acute infection (**Supplementary Table 1**). To better quantify the range of symptom severity within this mild cohort, a novel score was calculated representing the weighted time duration of symptomatic illness severity (see **Methods**). This illness severity score demonstrated the substantial clinical heterogeneity among participants (range 3-141; median 12) (**Fig. 1D**) and was used for further analyses. All COVID-19 participants were symptomatic during acute infection, but the most prevalent symptoms were fatigue or malaise (14/18), dry cough (13/18), headache (12/18), and loss of smell (11/18) (**Fig. 1D**). Most participants resolved symptoms within the acute period, but five female participants had persistent symptoms and were diagnosed with PASC.

The majority of participants mounted measurable SARS-CoV-2-specific adaptive immune responses. Antigen-specific CD4+ and CD8+ T cell responses targeting envelope (E), membrane (M), N, S, and open reading frame (ORF) 3, 6a, 7a, 7b, and 8 proteins peaked at approximately day 30 and did not appreciably decline in most participants (**Fig. 1E**). Frequencies of SARS-CoV-2-specific CD4+ T cells were consistently higher than CD8+ T cells within the same participant. Antibody responses were also analyzed for all participants, including neutralizing antibody and RBD-specific IgG titers (**Fig. 1F**): RBD-specific IgG titers and neutralizing antibodies peaked around day 30. While 100% (18/18) of COVID-19 participants mounted a sustained RBD-specific IgG response, only ~70% (13/18) sustained neutralizing antibody responses after acute infection. Spike-specific plasmablasts and IgG+ memory B cells were also detected in most participants (**Fig. 1G**). Peak plasmablast responses occurred within the first 15 days after onset of symptoms followed by a rapid decline by 30 days PSO in most participants. Spike-specific class-switched IgG+ memory B cells were also detected in all participants and generally increased in frequency over time. Antibody titers, and frequencies of CoV-2-reactive B and T cells were fit to linear and non-linear mixed-effects models using data >30 days PSO to estimate convalescent SARS-CoV-2 adaptive responses for each participant.

Next, we performed pairwise correlation analysis to identify connections among and between convalescent SARS-CoV-2-specific immune responses, demographic features, and clinical disease course (**Fig. 1H**). PASC was positively correlated with illness severity score, and an increased number and variety of symptoms (fatigue or malaise, headache, nausea or vomiting, and sore throat). Among immune correlates with PASC, the proportion of RBD-specific cells of total memory B cells was the strongest negative correlate suggesting potential association between PASC and reduced or delayed SARS-CoV-2 immune responses (see **Fig. 4**). Age was positively correlated with N IgG, and antigen-specific CD4+ and CD8+ T cell responses as seen in analysis of the broader cohort (Cohen et al., 2021). These results led us to perform an additional exploration of age-associated effects in this mild COVID-19 cohort and to perform a deeper analysis of participants with PASC.

### Early acute SARS-CoV-2 infection is characterized by an activated, inflammatory state including age-enhanced IFN responses and plasmablast expansion

We sought to define the immune response in mild COVID-19 during early acute infection, here ≤15 days PSO. Global unsupervised principal component analysis of the serum proteome showed significant differences between COVID-19 participants and uninfected controls (**Fig. S1A**). We compared early acute COVID-19 infection to uninfected controls and identified 407 differentially expressed proteins (**Fig. 2A; Supplementary Table 2A**). The serum proteome suggests a coordinated innate immune response that is dominated by type I IFN-related antiviral responses and proinflammatory cytokines (tumor necrosis factor, TNF; IL-18). The most differentially increased protein was RIG-I, an innate sensor encoded by the gene *DDX58* that detects double-stranded RNA viruses and drives expression of type I IFNs. RIG-I is known to restrict SARS-CoV-2 infection of pulmonary epithelial cells via type I/III IFN responses (Yamada et al., 2021) and early treatment with topical, nebulized IFNα2b resulted in reduced hospitalization and mortality (Wang et al., 2020b). FKBP5, a product of the acute stress response, was also differentially increased and was recently shown to regulate RIG-I signaling in response to influenza infection (Hao et al., 2020a). Upregulation of SAMD9L, a host restriction factor for poxviruses and an ISG, is also preferentially induced by type I IFNs (Pappas et al., 2009). Increased IFNλ1 suggested type III IFN responses were also induced. Many of the proteins that were differentially increased overlap with the serum proteome reported in hospitalized and severe COVID-19 (Filbin et al., 2020). In this previous study, strikingly, severe participants demonstrated a pronounced increase in IFNγ expression and increased type II IFN-driven protein expression. In contrast, IFNγ was not significantly elevated in our cohort, suggesting a less pronounced type II IFN-driven response may distinguish mild from severe COVID-19.

**Fig. 2:**
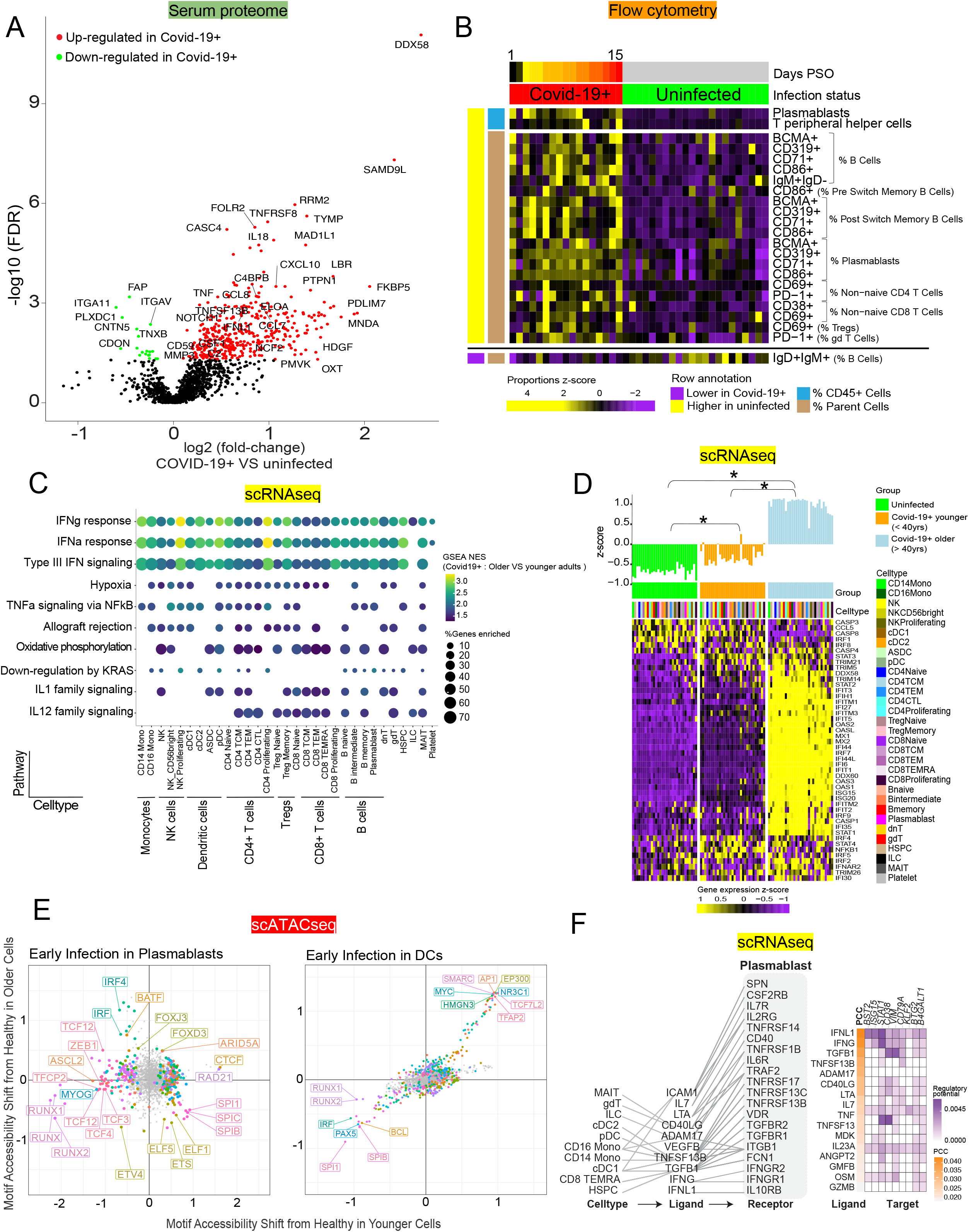
Early acute infection in mild COVID-19 is characterized by activation of T and B cells, and age-enhanced signatures of interferon signaling. **A)** Volcano plot showing differential expression of serum proteins between COVID-19 participants at ≤ 15 days PSO (n=18) compared to uninfected controls (n=22). Significantly differential proteins (FDR < 0.05) are colored as upregulated (red) or downregulated (green). **B)** Flow cytometry analysis shows cell type proportion differences between COVID-19 participants (n=18) compared to uninfected controls (n=23). Parent populations are plotted as % of CD45+ PBMCs, while phenotypic markers are plotted as % of parent. Differential populations were determined by fitting a linear model with change in proportions as a function of infection status adjusted for age and sex. Only significant proportions with adjusted p-values < 0.05 for infection status are shown. **C)** GSEA performed on scRNAseq using differentially expressed genes from comparison of older COVID-19 participants to younger COVID-19 participants in early acute infection within each cell type. Size of dots represents the percent of genes in each pathway that were enriched. Dot color represents the normalized enrichment score (NES) comparing older COVID-19 participants to younger COVID-19 participants. The top 10 most significant pathway enrichments are shown. **D)** Expression of genes enriched in IFN response pathways per cell type grouped by participant age and infection status. The column annotation bars represent the average gene expression across single cells per cell type and are scaled across samples. Gene expression was compared between groups using a Wilcoxon rank-sum test; asterisks indicate p < 0.05. **E)** Transcription factor motif analysis on scATACseq data showing differential accessibility of age-matched COVID-19 participants compared to uninfected controls in plasmablasts. Significantly enriched motifs were colored by motif family (Wilcoxon, FDR < 0.1), and the motifs with the largest shifts in each age subgroup (younger and older participants) are labeled. **F)** Predicted ligands with signaling activity in plasmablasts during early acute infection. Ligand-receptor interaction analysis was performed using DEGs from scRNAseq as input for NicheNet. The top 10 predicted ligands are shown ranked by Pearson’s correlation coefficient (PCC) of target genes. Potential source cell types were identified among PBMCs based on expression of the predicted ligands.

Cellular immune changes were assessed during acute infection by first analyzing proportions of innate and adaptive immune cell types and expression of phenotypic surface markers by flow cytometry. Principal component analysis of cell type frequencies showed significant differences between COVID-19 participants and uninfected controls (**Fig. S1B-C**). Bulk plasmablasts and PD-1^high^ CXCR5-T peripheral helper (Tph) cell frequencies were most significantly increased in COVID-19, while naive (IgD+IgM+) B cells were significantly decreased (**Fig. 2B)**. Activation marker-positive bulk B and T cells were also significantly increased, including CD69 and PD-1 on non-naive CD4+ T cells, CD69 and CD38 on CD8+ T cells, and BCMA, CD71, CD86, and CD319 (SLAMF7) on bulk plasmablasts and B cells. These changes demonstrate coordinated immune activation across adaptive and innate immunity in acute infection. scRNAseq provided deeper phenotyping of transcriptional state to analyze cell type activation. Cells were labeled by mapping scRNAseq to a CITE-seq-based multimodal reference atlas (Hao et al., 2020b), and resulting proportions were significantly correlated with flow cytometry (**Fig. S1D-H**). Bulk proliferating lymphocytes (CD71+ CD4+ and CD8+ T cells, CD71+ NK cells; **Fig. S2A**) were also significantly upregulated in mild COVID-19 infection, consistent with previous reports of a proliferating T cell phenotype in more severe COVID-19 (Stephenson et al., 2021).

Age is a well-known risk factor in COVID-19 that is associated with many compositional and functional changes (Channappanavar and Perlman, 2020; Richardson et al., 2020; Takahashi et al., 2020). Older COVID-19 participants (≥ 40 years old) had higher proportions of innate and adaptive immune cell types during early acute infection, including central memory CD4+ T cells (TCM), memory Tregs and CD14+ monocytes, while younger participants (<40 years old) had higher naive CD8+ T cells and naive Tregs (**Fig. S2B**). Serum levels of the inflammatory cytokine IL-6 and chemokines (CCL8, CXCL10) were significantly higher in older participants compared to younger participants during early acute infection (**Fig. S2C**). Transcriptomic analysis of cells by scRNAseq showed robust IFN responses in all cell types from older COVID-19 participants (**Fig. S2D-E, Supplementary Table 2B**) compared to uninfected controls. In contrast, only CD14+ monocytes and CD56^bright^ NK cells showed significant IFN responses in younger COVID-19 participants (**Fig. S2E**). Comparing the fold-change of these responses between older and younger COVID-19 participants revealed that IFN responses were significantly higher in all analyzed cell types from older participants (**Fig. 2C, Fig. S2F, Supplementary Table 2C**), with the greatest age-associated enhancement in the monocyte and NK populations. Differentially expressed ISGs showed a gradient of expression that increased with age and did not vary across cell types: uninfected participants had the lowest ISG expression, followed by higher expression in younger COVID-19 participants, and highest expression in older COVID-19 participants (**Fig. 2D**). While IFN responses were significantly activated in diverse PBMCs as determined by gene expression, only IFNλ1 (type III) was elevated in COVID-19 serum compared to uninfected controls in early acute infection (**Fig. 2A**). Other inflammatory cytokine signaling pathways were also up-regulated in diverse cell types from older participants (**Fig. 2C**). The TNF response was most prevalent following IFN responses, with age-enhanced responses in monocytes, CD56^bright^ and proliferating NK cells, DCs, CD4+ T cells, and B cells. IL-1 signaling was also significantly up-regulated in cell types from older participants, which may reflect inflammasome activation in early acute infection.

Transcription factor motif analysis on scATACseq was used to identify the most active cell types in acute infection, and any differences associated with age. Plasmablasts and innate immune cells (DCs, monocytes) had the largest motif shifts from uninfected controls (**Fig. S2G**). AP-1, a regulator of stress response and immune activation, was significantly enriched in innate immune cells, particularly CD14+ and CD16+ monocytes in older and younger participants, and DCs in older participants (**Fig. S2H**). These observations are consistent with scRNAseq and proteome data showing increased inflammatory cytokine responses that could activate AP-1. Enrichment of IRF motifs was also observed, particularly in plasmablasts, supporting strong IFN responses from transcriptional data.

One of the most striking differences in early infection was the increased abundance of circulating plasmablasts, which has typically been reported in more severe COVID-19 (Arunachalam et al., 2020; Kuri-Cervantes et al., 2020; Mathew et al., 2020). Bulk plasmablast proportions by flow cytometry and scRNAseq were significantly higher in COVID-19 participants ≤15 days PSO when compared to uninfected controls (**Fig. 2B, Fig. S2A**). Our data demonstrated that a high portion of the plasmablast expansion in mild SARS-CoV-2 was S-specific (**Fig. 1G**). Plasmablasts from younger COVID-19 participants compared to uninfected controls were enriched for RAD12 and CTCF motifs, and were strongly depleted for TCF family and RUNX motifs. Consistent with age-enhanced IFN responses, we found IRF family motifs were significantly enriched in plasmablasts from older participants (**Fig. 2E**, left panel). In contrast to plasmablasts, DCs had the second highest magnitude change in motifs, but showed few age-associated changes (**Fig. 2E**, right panel): all participants showed enrichment of AP-1 and high-mobility group (HMG) family members HMGN2 and TCF7L2, and depletion of RUNX and SPI family motifs. These results suggest a potential link between IFN and plasmablast responses during acute infection by SARS-CoV-2.

We inferred ligand-receptor (LR) interactions from plasmablast transcriptomes during early acute infection to define the upstream signals that may drive their response (Browaeys et al., 2020) (**Fig. 2F**). The top ligands identified as inputs into early plasmablasts included type II and III IFNs (IFNγ, IFNλ1), TNFRSF13B/TACI, and CD40LG (AUROC>0.58). Potential PBMC sources were identified including innate immune cells for TNFSF13B, and MAIT and γδ T cells for CD40LG. IFNγ and IFNλ1 had no putative PBMC sources identified, suggesting tissues may be a primary source for IFNs in early acute infection. STAT1 was identified as a shared downstream target for these ligand-receptor interactions (**Fig. 2F**; **Supplementary Table 2D**), consistent with strong IFN responses. These results support the generation of mechanistic hypotheses for key signals identified in other studies, such as the role of early IFNL1 in mild COVID-19 (Galani et al., 2021). Our data suggest early IFN, particularly IFNλ1, may drive plasmablast responses, potentially through synergy with BCR (Syedbasha et al., 2020) and Toll-like receptor (TLR) signaling (de Groen et al., 2015), and this may link early control of viral replication, inflammatory cytokine milieu, and extrafollicular B cell responses. Future studies will be critical to confirming the function of these plasmablasts in SARS-CoV-2 infection.

### Longitudinal analysis shows that a decrease in activated and proinflammatory responses correlated with an increase in repair and convalescent homeostasis pathways

After defining unique characteristics of early acute infection, we analyzed significant longitudinal changes in immune phenotype during mild COVID-19. Serum proteins that significantly decreased over time were largely proinflammatory proteins, including cytokines (IL-18, TNF), chemokines (CCL7, CCL8, CXCL10), complement proteins (C4BPB), and IFNλ1 (**Fig. S3A**). These proteins were enriched for type I and II IFN responses, TNF signaling, interleukin (IL-2, IL-6) signaling, and innate immune sensor (TLR; Nod-like receptor, NLR) signaling (**Fig. 3A, Supplementary Table 3A**). Inflammatory protein changes detected during acute infection resolved around day 30 PSO for most COVID-19 participants. A subset of proteins increased over time, and these were enriched for epithelial-mesenchymal transition (EMT), angiogenesis, and coagulation pathways. These changes suggest that convalescence is characterized by a transition to a wound healing, homeostatic phenotype from the inflammatory response in acute infection (**Fig. 3A**). Some chemokines increased over time, including CCL16, an IL-10 inducible chemokine that can serve both proinflammatory and immunoregulatory functions, and is consistent with previous observations of CCL16 depletion during acute infection in severe COVID-19 (Filbin et al., 2020). Notably, some inflammatory responses persisted in PASC, particularly for the sickest participants (orange and red PASC column annotation, respectively; **Fig. 3A**).

**Fig. 3:**
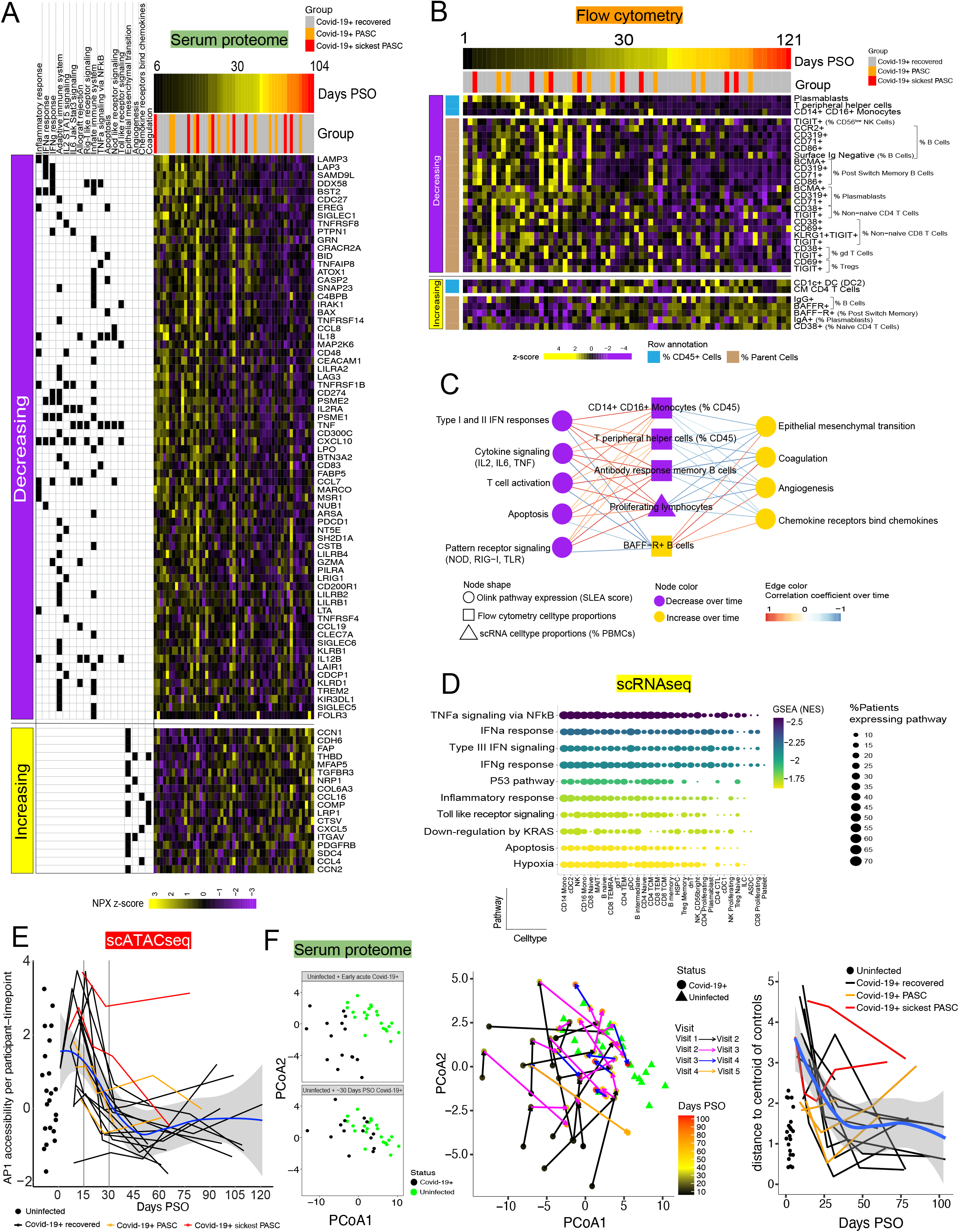
Longitudinal analysis shows individual heterogeneity in path to convalescence with resolution of inflammatory cytokine signaling coordinated with increasing wound healing and EMT. **A)** Heatmap of changes in serum proteins over time in mild COVID participants. Proteins that significantly changed over time were identified and enriched for pathways using Fisher’s overlap test. Individual columns represent samples ordered by days PSO, with column annotations for days PSO, and recovery status as either PASC (orange), sickest PASC (red), or recovered (grey). Rows represent individual proteins. Features were considered significant with adjusted p-values < 0.05. **B)** Heatmap of longitudinally changing cell frequencies quantified by flow cytometry among COVID-19 participants. Columns represent individual samples ordered by days PSO, with column annotations for days PSO, and recovery status as either PASC (orange), sickest PASC (red), or recovered (grey). Rows represent significant cell frequencies shown as row-scaled % of total CD45+ cells or % of parent population. **C)** Pairwise correlations between functional pathways (circles, serum proteome) and cell frequencies (squares, flow cytometry; triangles, scRNAseq) over time. The “antibody response memory B cells” node combines plasmablasts (% of CD45+ cells), and BCMA+, CD71+, and CD86+ (% of post-switch memory B cells) as determined by flow cytometry. The “proliferating lymphocytes” node constitutes CD4, CD8, and NK proliferating cells (% of PBMCs) as obtained by scRNAseq. The “BAFF-R+ B cells” node constitutes BAFF-R+ (% B cells) and BAFF-R+ (% of post-switch memory B cells). The edges to these specific nodes represent the average Spearman correlation coefficient across these cell types to individual serum proteome features. Edges are colored by magnitude of positive (red) or negative (blue) Spearman correlation coefficients. Only significant correlations (adjusted p-values < 0.05) are shown. **D)** GSEA performed on genes from scRNAseq that were longitudinally decreasing over time per participant by cell type. Dot size represents the percent of COVID-19 participants that showed enrichment for the indicated pathway and cell type. Dot color represents the negative NES indicating the magnitude of decreasing pathway expression over time. The top 10 significant pathway enrichments are shown. **E)** Longitudinal transcription factor motif accessibility for AP-1 in CD14+ monocytes from COVID-19 participants. Uninfected participants are represented by black solid circles on the left. AP-1 significantly decreased over time from day 10 onwards in the overall cohort (p-value < 0.0001). **F)** Principal coordinates analysis on all longitudinally changing proteins in COVID-19 participants was used to visualize heterogeneity in the path to convalescence. Early acute COVID-19 samples (***left top***) and ~30 days PSO (***left bottom***) were plotted with uninfected controls. The **middle** panel shows the progression of COVID-19 participants from early (black symbols) to late (red symbols) days PSO. Most COVID-19 participants (solid circles) were well separated from uninfected participants (green triangles) during acute infection, but converged to the uninfected region as they recovered. The right panel shows the corresponding distances to the centroid of uninfected participants (solid black circles). The distances of uninfected participants were plotted with solid circles and randomly assigned days PSO between 0 and 5. The distances of recovered COVID-19 participants were plotted with black solid lines, while those of PASC and sickest PASC participants were plotted with orange and red solid lines, respectively. Loess smoothing (blue line with 95% confidence intervals shaded grey) was used to guide the eye.

Cell proportions measured by flow cytometry mirrored the proteome, with most changes during acute infection resolving by 30 days PSO. These included decreases in bulk plasmablasts, Tph cells, and CD14+ CD16+ intermediate monocytes, and activation marker-positive B and T cells (**Fig. 3B**). CD1c+ DC2s and CD4+ TCMs increased over time, consistent with a return from virus-infected tissues for DC2s (Bosteels et al., 2020; Gill et al., 2008) and expansion or differentiation of CD4+ TCM cells after acute infection. BAFF-R+ class-switched memory B cells and IgG+ B cells increased over time, which suggests recent memory differentiation and class-switching from activated B cells during acute infection (Lau et al., 2021). IgA+ plasmablasts also increased over time and may be a consequence of acute infection-induced plasmablasts contracting. Bulk proliferating lymphocytes identified in scRNAseq also significantly declined over time (**Fig. S3B**).

To assess coordination of the immune response, we correlated the change in cell type proportions measured by flow cytometry and scRNAseq with changes in the serum proteome over time (**Fig. S3C**). Cell types that positively correlated with the activity of the most serum proteome pathways included CD14+ CD16+ monocytes, Tph, memory B cells, and proliferating lymphocytes (**Fig. 3C**). Decreases in these cell type proportions over time were associated with decreased type I and II IFN responses, inflammatory cytokine and immune activation signaling, and PRR signaling. In contrast, these decreases were correlated with increases in BAFF-R+ B cells and serum proteome pathways for chemokines, wound healing, and tissue repair, suggesting temporal coordination between inflammatory resolution and homeostatic repair characterizes successful recovery from mild COVID-19.

We next used scRNAseq and scATACseq data to longitudinally map cell type-specific changes in function over time. Genes demonstrating significant changes over time were identified per cell type, and these changing genes were then mapped to their corresponding pathways. Pathways that showed the top enrichment scores and decreased the most over time, as weighted by number of impacted cell types and participants (see Methods), included TNF signaling, IFN responses, TLR signaling, and apoptosis (**Fig. 3D, Fig. S3D, Supplementary table 3B**). These pathways were consistent with inflammatory protein decreases in serum. Innate immune cells (monocytes, cDC2s, NK subsets) represented the majority of the top cell types expressing these longitudinally changing pathways. scATACseq showed that most transcriptional motifs enriched in acute infection rapidly declined in accessibility over time, as demonstrated by IRF family motif shifts (**Fig S3E**). AP-1 motifs were among the most accessible features in innate immune cell types during early acute infection for nearly all COVID-19 participants (**Fig. S2H**). Accessibility of AP-1 motifs peaked in CD14+ monocytes during acute infection, followed by a rapid decline to homeostatic baseline by 30 days PSO (**Fig. 3E**). PASC participants, especially the sickest PASC, were an exception, maintaining higher accessibility >30 days PSO, which may represent persistent innate immune activation. Principal coordinates analysis (PCoA) on longitudinally changing immune features revealed the individual trajectories of COVID-19 participants from acute infection through convalescence (**Fig. 3F**). Early acute infection samples (black circles) showed clear differences from uninfected controls (light green circles, **Fig. 3F**, top inset left), while most participants around 30 days PSO were closer to uninfected controls (**Fig. 3F**, bottom inset left). COVID-19 participant trajectories (lighter colored circles and arrows) converged closer to the uninfected control space by ~30 days PSO (**Fig. 3F**, middle panel). Distances for each sample were calculated from the centroid of the uninfected control group to visualize the timing of inflammatory resolution (**Fig. 3F**, right panel). Most participants’ trajectories resolved near uninfected controls around day 30, with notable exceptions for the two sickest PASC participants who remained >2.5 distances from the uninfected group. Overlaying comorbidity annotations on the PCoA revealed that the sickest PASC participant with the most comorbidities had the most distinct trajectory from other COVID-19 participants (**Fig. S3F**). Altogether, longitudinal analyses across data types demonstrated that most mild COVID-19 participants resolved into a convalescent immune phenotype similar to uninfected controls, while PASC participants have persistent differences and unique trajectories.

### PASC participants are distinguished by dampened anti-viral and IFN responses in acute infection and persistent, unresolved inflammatory signaling in convalescence

We sought to perform deeper immune profiling of PASC participants compared to recovered COVID-19 participants. Recovered COVID-19 participants had no symptoms after ~2 weeks PSO (range: 0-17 days PSO) However, the five participants who were classified as PASC had continued symptoms lasting for > 60 days PSO. All PASC participants were female, similar to the sex bias reported in previous studies (Davis et al., 2020; Evans et al., 2021; Sigfrid et al., 2021). Two participants who had more severe illness and persistent symptoms were categorized as sickest PASC due to the significant impact of ongoing symptoms on activities of daily living, quality of life and ability to continue employment. These symptoms included neurological symptoms, alterations in smell and taste, chest tightness and joint pain (PTID 795172; illness severity score 71, WHO score 3), and cardiovascular abnormalities, headache, fatigue, and change in taste (PTID 523731; illness severity score 141, WHO score 3) (**Supplementary Table 1A**). PASC participants had qualitative differences in CoV-2-specific responses (**Fig. 4A**), but none were statistically significant, likely due to the small sample size. There are however trends suggesting that PASC participants may have dysregulated adaptive immune responses. Most notably, the two sickest PASC participants had low or absent SARS-CoV-2-specific CD4+ and CD8+ T cell responses after 30 days PSO, high RBD IgA, and low RBD IgM. PASC participants also had SARS-CoV-2 response estimates below the lower quartile of recovered COVID-19 participants RBD- and S-specific memory B cells, and N IgG (**Fig. 4A**).

**Fig. 4:**
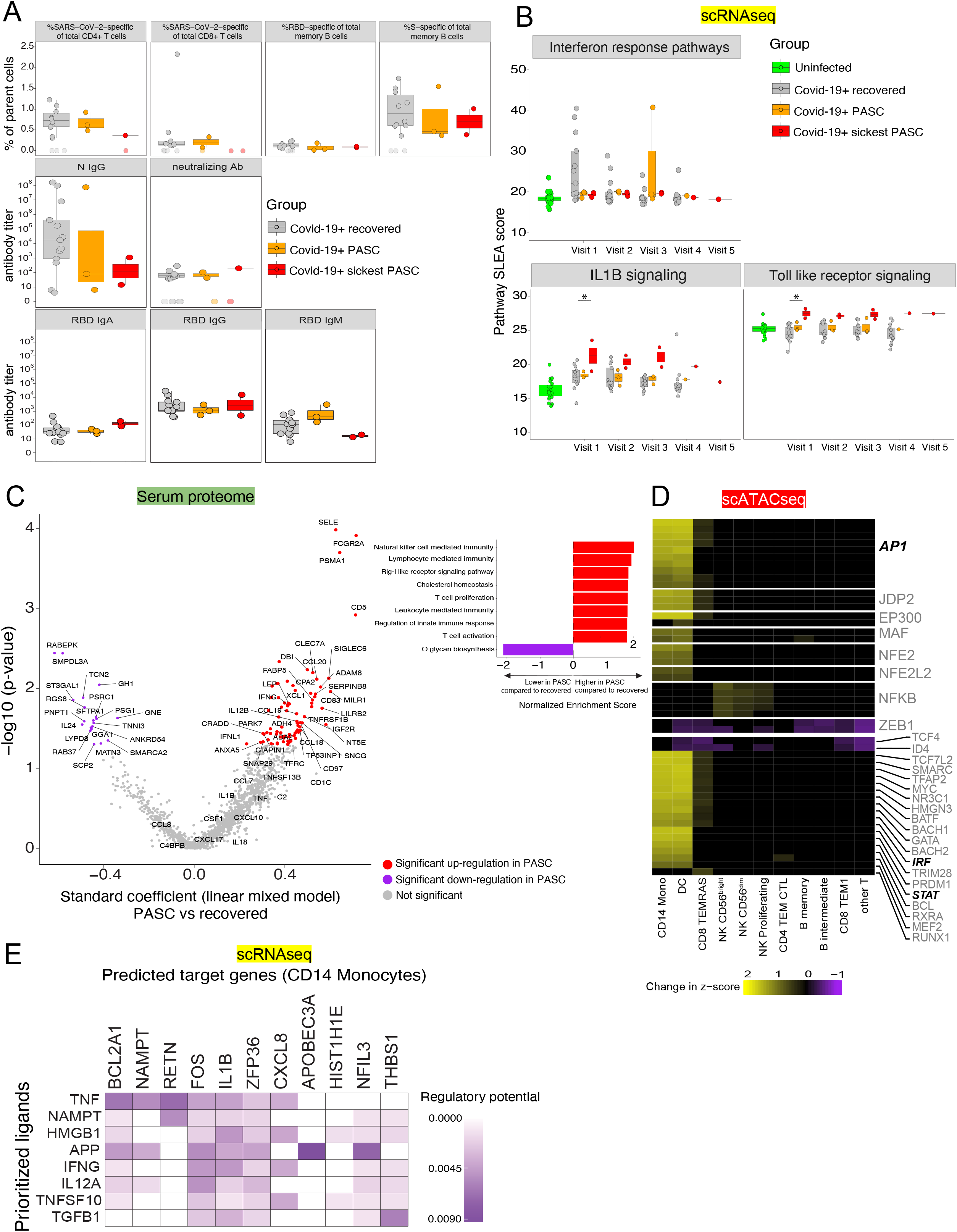
Persistent immune activation distinguishes post-acute sequelae of SARS-CoV2 infection (PASC) from recovered COVID-19. **A)** SARS-CoV-2-specific adaptive immune responses estimated at day 30, except neutralizing antibodies at day 42, and comparing PASC and recovered COVID-19 participants. Each symbol represents 1 participant. Undetectable estimates are denoted by empty symbols. **B)** scRNAseq SLEA pathway scores of the significantly different pathways in CD14+ monocytes. GSEA was performed comparing all COVID-19 participants in early acute infection to uninfected controls. The p-values of GSEA were <0.05. The SLEA pathway score per sample was then calculated and tracked over time. **C)** Volcano plot (left panel) of the differentially expressed serum proteins over time comparing PASC to recovered COVID-19 participants. Each protein’s expression was fitted to a linear mixed-effects model. Significantly differential proteins are shown with p-values < 0.05 are shown in red and purple dots. Results of the Gene Set Enrichment Analysis among all differential proteins are represented in the bar plot on the upper right panel, with the top five pathways (ranked by the NES) up-regulated (red) and down-regulated (purple) in PASC compared to recovered participants. Part of the proteins enriched among these pathways are also shown in the volcano plot. **D)** Transcription factor motif analysis of scATACseq data in samples >30 days PSO. PASC participants were compared to recovered COVID-19 participants and enrichments were calculated using ChromVar motif z-scores. The top 50 largest significant motif differences are shown (Wilcoxon FDR < 0.10). **E)** Ligand-receptor interaction analysis was used to identify predicted signals driving innate (CD14+ monocyte) cells in PASC participants. The top 5 predicted ligands and their target gene signatures are shown. Ligands were ranked by Pearson correlation coefficient between expected and observed target gene expression.

To identify immune pathways and cell types that define PASC participants, we evaluated scRNAseq data from PASC compared to those who had an uneventful recovery from acute SARS-CoV-2 infection. Differentially expressed genes (DEGs) were identified by comparing all PASC participants to recovered COVID-19 participants ≤15 days PSO by cell type. CD14+ monocytes, CD8+ TEMRAs and CD8+ effector memory T cells (CD8+ TEM) showed the highest number of DEGs (**Fig. S4A**). Mapping DEGs to pathways by cell type revealed dysregulated signaling in early acute infection that persisted longitudinally in PASC. Consistent with evidence of dysregulated immune responses described above, early infection in PASC was characterized by significantly lower IFN responses compared to recovered participants in CD14+ monocytes, CD8+ TEMRAs and CD8+ TEM cells (**Fig. 4B upper panel, Fig. S4B**). These signatures remained persistently low throughout disease, while expression levels in recovered participants were higher during acute infection and declined over time. Recovered participants showed coordinated expression of genes from these pathways, which define an antiviral response (**Fig. S4C, Supplementary Table 4A)**. CD14+ monocytes from PASC participants also demonstrated significantly higher expression of proinflammatory pathways (IL-1β signaling, TLR signaling) in early acute infection compared to recovered participants, with the highest responses observed in the sickest PASC participants and persisting throughout infection as well as the post-acute period (**Fig 4B, lower panels**). The up-regulated gene signature included *IL1B*, *CXCL8, HIF1A, S100 family genes* (*S100A8, S100A9, S100A12*), TLRs (*TLR2, TLR4, TLR8*), and the AP-1 transcription factor subunit (*FOS*) (**Fig. S4D, Supplementary Table 4A**). In contrast, IFN responses in CD14+ monocytes were lower in PASC participants during early acute infection compared to recovered, but these responses also persisted in PASC participants after 30 days PSO (**Fig. S4E**). These results suggested PASC is characterized by persistent inflammation compared to timely resolution in successful recovery. In addition, both CD8+ TEMRAs and TEMs had significantly higher gene expression of TNFα signaling via NFκB in PASC compared to recovered participants during early acute infection, with highest elevation in sickest PASC, and this persisted throughout infection (**Fig. S4F**).

The presence of an inflammatory gene signature led us to examine whether an inflammatory protein signature also defined PASC. We analyzed the longitudinal proteomes of an expanded PASC cohort including 60 PASC participants and 26 recovered COVID-19 participants. Outlier analysis on the longitudinal serum proteome identified protein signatures that distinguished PASC from recovered COVID-19 participants. Most recovered COVID-19 participants had a decreasing number of differential proteins after early acute infection compared to uninfected controls indicating their path back to normalcy (**Fig. S4G**). In contrast, PASC participants had stable or increasing numbers of differentially expressed proteins over time. PASC participants were defined by a persistently elevated inflammatory signature throughout the course of infection. These included significantly up-regulated GO terms like NK cell mediated immunity, RIG-I-like receptor signaling, T cell activation, T cell proliferation, and regulation of innate immune repsonse among others, which were enriched for proteins such as IFNγ, IFNλ1, IL-1β, IL-12, IL-18, and TNF (**Fig. 4C, Supplementary Table 4B**). Significantly upregulated proteins in PASC also included members of the TNF superfamily in PASC, including TNFSF13B/soluble BAFF receptor and TNFRSF1B/TNF receptor 2, consistent with immune activation (**Supplementary Table 4B**). Overall, analysis of the serum proteome of PASC participants provides further evidence of a hyperinflammatory state.

In an effort to understand whether there are particular immune cells either driving or responding to the signals identified in the serum proteomic studies, we performed transcription factor (TF) motif analysis of scATACseq data. This revealed key TF motifs that correlate with aberrant cell phenotypes in PASC participants compared to COVID-19 recovered participants at >30 days PSO. A set of AP-1 family motifs were significantly enriched in dendritic cells (DCs) and CD14+ monocytes. Paired with evidence of increased expression of the genes encoding the AP-1 subunits *FOS* and *JUN*, this suggests a state of persistent immune activation that is most prominent in innate myeloid cells (**Fig. 4D**). Other prominent enriched motifs included BACH, IRF, and STAT families, all associated with persistent inflammatory cytokine signaling. The same motifs were also enriched, albeit to a lesser extent, in CD8+ TEMRAs. NK cells specifically showed enrichment of NF-κB motifs. Motif enrichments were not observed across all innate immune cells or for most adaptive immune cell types in PASC. Using scATACseq, we identified genes nearest to differentially accessible PASC peaks with AP-1 motifs and analyzed their expression using scRNAseq data to compare PASC to recovered participants after 30 days PSO. Multiple genes were significantly up-regulated in PASC and these were enriched for inflammatory pathways including TNF signaling, IL1B signaling, TLR signaling, IFN responses, and hypoxia (**Fig. S4H**). This agreement between scATACseq and scRNAseq suggests that inflammatory signals activating AP-1 contribute to persistent inflammation in PASC.

To identify potential signals driving the phenotype of CD14+ monocytes in female PASC participants, scRNAseq gene expression changes were used to predict ligand-receptor interactions by the NicheNet method (Browaeys et al., 2020). Ligand activity was predicted and ranked by the correlation between knowledge-based predictions and experimentally observed levels of target gene expression (**Fig. S4I**). Ligand expression was assessed in all cell types along with the fold increase in their expression per cell type compared to recovered participants to identify PBMC sources of ligand signals (**Fig. S4J)**. TNF, NAMPT and IFNG were among the top predicted ligands with highest target gene activity in CD14+ monocytes (**Fig. S4I, S4J, 4E**). Diverse potential sources for these ligands among PBMCs included CD14+ monocytes, dendritic cells, proliferating CD8+/CD4+ T cells and proliferating NK cells (**Fig. S4J)**. Target genes of these predicted ligands included *IL1B*, *FOS*, *NAMPT*, *BCL2A1*, and *CXCL8* (**Fig. 4E, S4K**), which confirm observations in **Fig 4B and Fig. S4D**. Overall, these data suggested a key role for the cytokine milieu driving a persistently proinflammatory state in CD14+ monocytes, identifying biomarkers for diagnosis of PASC, and revealing multiple therapeutic targets.

These results suggest that PASC is characterized by blunted innate and adaptive antiviral immune responses during acute infection that leads to a persistent inflammatory state including increased expression of proinflammatory genes and proteins. DCs and monocytes demonstrated the most significant increases in chromatin accessibility including motifs for AP-1, STAT, and IRF family transcription factors known to drive inflammatory gene expression. There is clear evidence of persistent inflammation in PASC, but It remains unknown whether these are the primary pathogenic mechanisms causing PASC or merely correlates.

### Heightened early acute interferon and antiviral signaling correlate with stronger humoral responses to SARS-CoV-2 in convalescence

Antibody responses are a key component to controlling viral infection, disease outcome, and protection from reinfection. Clinical trials have directly shown that neutralizing antibodies (nAb) to SARS-CoV-2 can reduce mortality and length of hospitalization (Chen et al., 2021). Immune responses in acute infection are critical for establishing the coordinated immune response required for development of antibody and memory B cell responses in convalescence. Serum protein expression and flow cytometry proportions were estimated at day 7 PSO. RBD IgG titers and S-specific memory B cell frequencies at day 30 PSO and neutralizing antibody (nAb) titers at day 42 PSO were estimated by linear mixed-effects models. Observed and estimated values were strongly correlated (**Fig. S5A, S5B**).

We identified a signature of circulating proteins, present during early acute infection (day 7 PSO), that strongly correlated to peak antibody titers for RBD IgG and neutralizing antibody (nAb) at days 30 and 42 respectively (**Fig. 5A, 5B**). A circulating protein signature was also identified that correlated with spike-specific memory B cell frequencies (**Fig S5C**). Correlations were also found between specific immune cell subsets identified by flow cytometry and RBD IgG, nAb, and the S-specific memory B cell frequency (**Fig. 5D, 5E; Fig. S5D**). Analysis of the serum protein milieu at day 7 PSO revealed that higher expression of proteins in inflammatory pathways were significantly correlated with higher RBD IgG titers at day 30 PSO. Among the most prominent were ISGs (TRIM21, SAMD9L), complement (CD55, GPB1A), and innate sensor signaling proteins (TANK, MAVS, RIG-I) (**Fig. 5A**). Neutralizing antibody titers at day 42 were also positively correlated with day 7 innate immune proteins (IFNλ1, DDX58), while negative correlates included a subset of inflammatory proteins (IL-34, IL-17A, CD209/DC-SIGN) and chemokines (CCL15, CCL28) (**Fig. 5B**). These underscore the importance of innate immunity in early infection for encoding the magnitude of the convalescent antibody response. Sixteen serum proteins were common correlates for both RBD IgG and nAb (**Fig. 5C**). Soluble correlates were complemented by positive correlations with flow cytometry populations, including selected activation marker-positive T cells (PD-1+/TIGIT+) and naive B cells (IgD+IgM-), as well as CD56^high^ NK cells (**Fig. 5D**). This links the magnitude of humoral response to the degree of T and B cell activation, potentially through support of B cell maturation to plasma and memory cells. Negative correlates of RBD IgG included pre-switch memory B cells (CD27+IgD+), and CD11b+ B cells. Cell proportions that positively correlated with nAbs included circulating bulk CXCR5+ PD-1+ Tfh cells, and CD56^high^ and CD16-NK cells. Phenotypic markers of activation on T cells (CD38, CD71, PD-1) and plasmablasts (BCMA) were positively correlated with neutralizing titers. Negative correlates of neutralizing titers included naive CD4+ T cells and inhibitory receptor-positive (KLRG1, TIGIT) CD56^low^ NK cells (**Fig. 5E**). CD56^high^ NK cells and CD38+ CD8+ T cells were shared positive correlates of both neutralizing antibodies and RBD IgG titers (**Fig. 5F**). Coordination of multiple aspects of immunity (lymphocyte activation, innate immune signaling) in early infection are strong predictors of effective priming for IgG and neutralizing antibody responses against SARS-CoV-2.

**Fig. 5:**
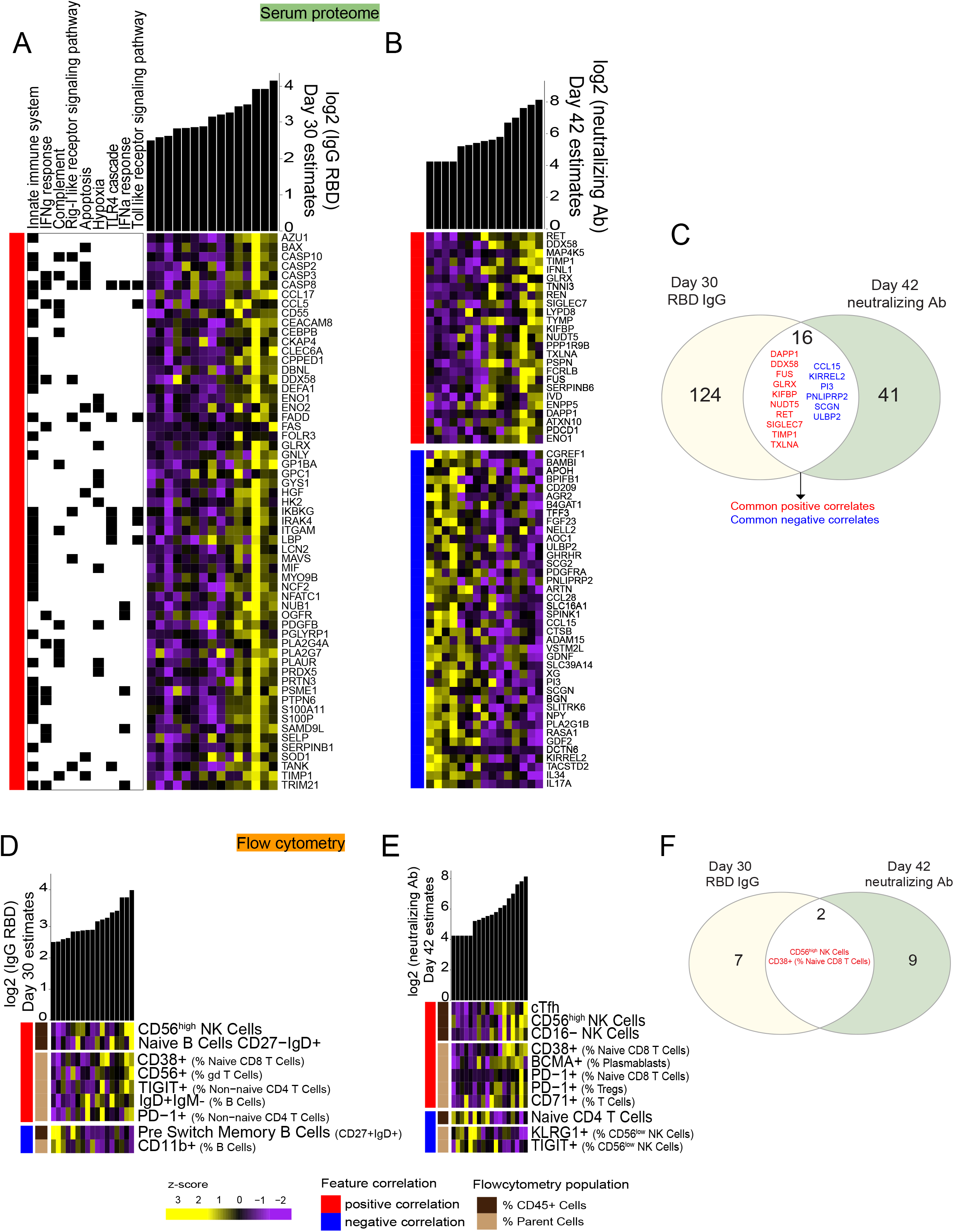
Immune correlates from early acute infection highlight the coordination of innate inflammatory response with humoral responses to SARS-CoV-2. Day 7 estimated serum proteins and pathway enrichments that significantly (Spearman’s correlation coefficients > 0.5 and adjusted p-values < 0.05) correlated with estimated day 30 RBD IgG titer (**A**) and day 42 neutralizing antibody titers (**B**). **C)** Venn diagram of the early acute protein feature overlaps that correlated with both day 30 RBD IgG and day 42 neutralizing antibody titers. Flow cytometry based cell proportions that significantly (Spearman’s correlation coefficients > 0.5 and adjusted p-values < 0.05) correlated with estimated day 30 RBD IgG titer (**D**) and day 42 neutralizing antibody titers (**E**). **F)** Venn diagram of the early acute cell proportion overlaps that correlated with both day 30 RBD IgG and day 42 neutralizing antibody titers. Rows correspond to proteins or cell frequencies measured by flow cytometry. Columns correspond to individual COVID-19 participants arranged by increasing order outcomes (represented as column annotation bars).

A critical role for memory B cells has been posited to provide rapid protection upon rechallenge (Dan et al., 2020). We identified serum proteins and cell frequencies from early infection that correlate with convalescent S-specific IgG+ memory B cell responses. Similar to IgG titers, we observed strong positive correlations with secreted proteins enriched in innate immunity, IFN responses, and RIG-I signaling (**Fig. S5C**). In contrast, proteins negatively correlated with memory B cells included complement, chemokines (CCL16, CCL20), and interleukin (IL-1, IL-2) signaling. Cell frequencies that positively correlated with memory B cell response included DC2s and CD25-ILCs, and activated plasmablasts and memory B cells (BCMA) (**Fig. S5D**). KLRG1 on non-naive CD4+ T cells and PD-L1 on CD14+CD16+ intermediate monocytes were also positively correlated with memory B cell responses. Overall, these findings identified diverse cellular and molecular features of early infection that correlate with antibody and memory B cell responses, identifying candidates for immunomonitoring and novel linkages between the acute antiviral response and the magnitude of humoral immune responses.

### Integrative analysis reveals key network nodes in acute SARS-CoV-2 infection, their longitudinal resolution, and persistence in PASC

We sought to trace the longitudinal resolution of immune signals that characterized acute infection through longitudinal resolution and convalescence. Several serum inflammatory proteins that were elevated during acute infection compared to uninfected controls resolved over time in recovered participants but showed prolonged elevation in PASC participants, including IFNγ, IFNλ1, IL-12p40, IL6, and TNF among others (**Fig. 6A, Fig. S6**). To infer the origin and signaling activity of these proteins, we performed intercellular communication analysis of scRNAseq data using NicheNet (**Methods**). We first identified cell type-specific expression of proteins elevated in early acute infection (from **Fig. 6A**), and subsequently determined candidate secreting/sender cell types for these signals (**Fig. S7A**) to define the ligand-cell type interaction network in PBMCs from acute infection (**Fig. 6B**, 46 predicted ligands in >10% of all cell types). Ten cell types were analyzed as potential receivers of signals based on changes observed in early acute infection (**Figs. 2, 3**): plasmablasts, CD14+ monocytes, CD16+ monocytes, NK cells, proliferating cells (CD4, CD8, NK), and DCs (DC1, DC2, pDC). The top 10 high-confidence predicted ligands among these receiver cell types were determined (**Fig. 6C**) and ranked based on the percent of cell types with predicted ligand activity. These ligands demonstrated the prevalence of inflammatory signaling among cell types during COVID-19 infection, including IFNγ, IFNλ1, IL-12, and TNF (**Supplementary Table 5A; Fig 6B-D**). Multiple downstream target genes were shared between the predicted ligands, including *STAT1* (**Fig. 6D**), *DDIT4* (regulator of mTOR, proliferation, autophagy), and other transcriptional regulators (*NFKBIA*, *JUN*, *FOS*).

**Fig. 6:**
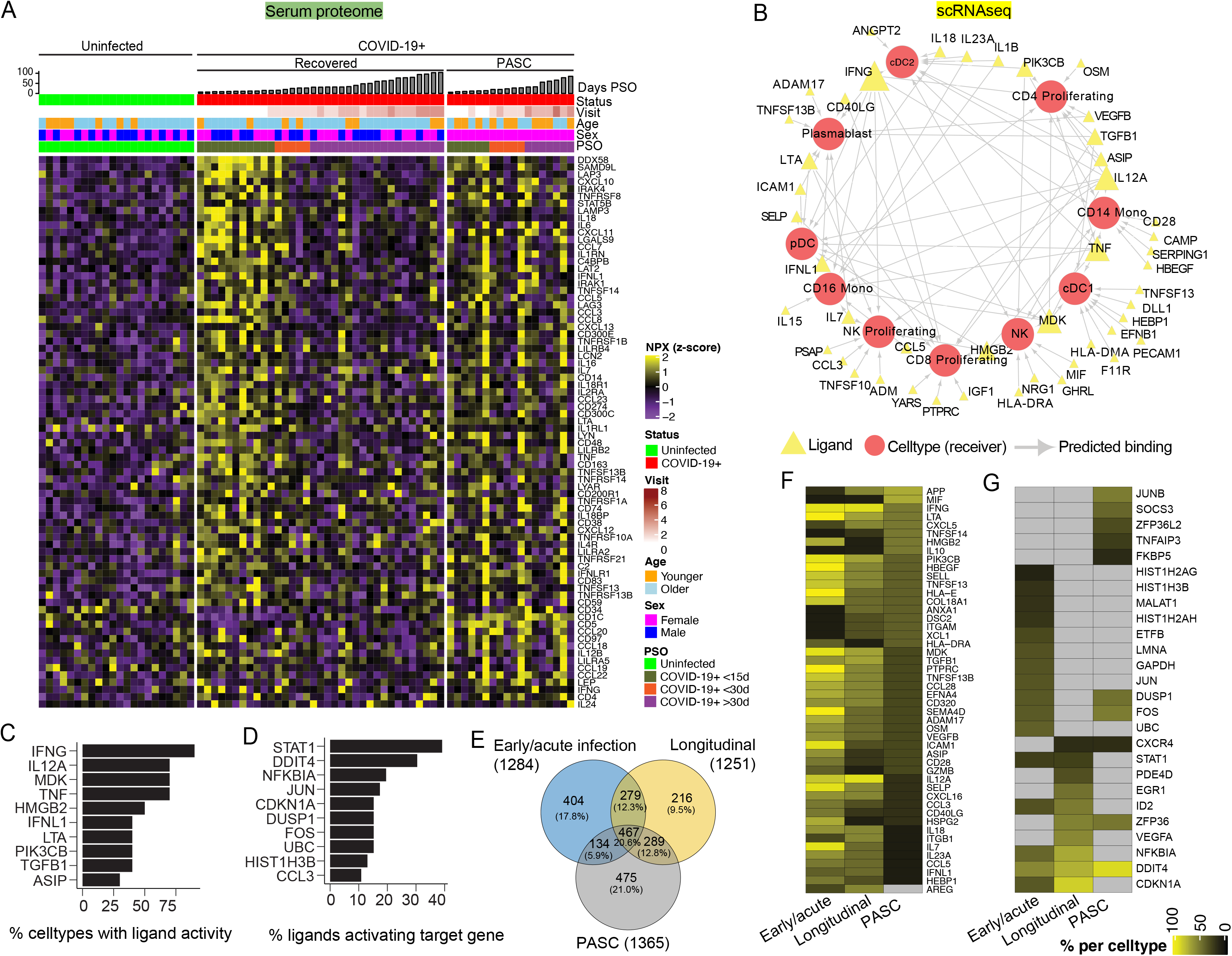
Integrative analysis uncovers key nodes for immunomonitoring and potential therapeutic targets throughout infection, convalescence, and PASC. **A)** Differentially expressed immune-related proteins observed in early acute infection (≤15 days PSO), longitudinally and in PASC COVID-19 participants derived from serum proteomics study. Among the broader serum proteomics cohort, only samples that had paired scRNAseq data are represented in this heatmap. **B)** NicheNet-based intercellular communication analyses of single cell RNA data from early acute COVID-19 infection participants. We retrieved top 10 inferred ligands influencing the ligand-target expression in receiver cell types. Triangles show the ligand and circles represent the receiver cell type. The edge between nodes and ligands shows the inferred relationship in early acute COVID-19 infection from scRNA data. The size of the triangle is proportional with the number of edges outgoing. **C)** The top 10 inferred ligands (per cell type) with the percent of cell types showing ligand activity, and **D)** the percentage of ligands activating downstream target genes in early acute COVID-19 infection were shown. **E)** The overlap between the inferred ligand-receptor interactions from differential intercellular communication between three subgroup comparisons: 1) early acute COVID-19 participants ≤ 15 days PSO compared to uninfected, 2) longitudinal COVID-19 at >15 days PSO compared to early acute COVID-19, and 3) PASC participants compared to recovered COVID-19 participants at all timepoints. Longitudinal changes in **F)** ligand and **G)** target usage in early acute COVID-19, longitudinal and PASC participants respectively are shown. Changes were calculated as the difference in predicted ligand per cell type **(F)** or downstream target genes regulated by predicted ligands **(G)** in the subgroup specified compared to respective control conditions listed above. Columns represent each subgroup and differences >10% were shown.

Next, we analyzed LR interactions of differentially expressed target genes from 3 distinct comparisons: 1) early acute infection compared to uninfected, 2) longitudinal timepoints (>15 days PSO) compared to early acute infection, and 3) PASC participants (all timepoints) compared to recovered COVID-19. There was significant overlap of LR pairs (467 pairs, ~21%) identified in early acute infection, longitudinal timepoints, and PASC participants (**Fig. 6E, Supplementary Table 5B**), indicating common signals driving each subset of mild COVID-19. PASC had the highest number of unique LR interactions (475 pairs, 21%) followed by early acute infection timepoints (404 pairs, ~18%), and longitudinal timepoints (216 pairs, ~10%). Notably, a subset of LR interactions (134, ~6%) were shared between early acute infection and PASC, which may represent signals from acute infection that persist abnormally in PASC. We dissected LR usage by participant subgroups to identify ligands and targets that were more abundantly used in PBMC networks for early acute infection, longitudinal resolution, and PASC (**Figs. 6F, G**). Multiple innate immune signals and inflammatory signaling were upregulated in early acute infection compared to uninfected controls, and persisted in PASC compared to recovered COVID-19 participants (**Fig. 6A, Fig. S6**). Increased activities of some predicted ligands in PASC participants were consistent with significantly elevated levels in the serum proteome, including IFNγ, IL-6, and IL-12 among others (**Fig. 6F, Supplementary Table 5C, Fig. S7B**). A subset of downstream gene targets from predicted ligands was used in at least two subgroups, and primarily consisted of genes coding intracellular proteins: *DDIT4, NFKBIA*, and *CDKN1A* (**Fig. 6G**).

Overall, data from multiple omics demonstrated that PASC participants showed enrichment for a subset of cytokines, chemokines, transcription factors (*STAT1*, AP-1 subunits *FOS* and *JUNB*), and genes proximal to these open transcription factor sites that were involved in inflammatory cascades such as TNF and IL-1β signaling pathways. The signaling landscape from PASC suggests neutralizing soluble cytokines, such as TNF or IFNγ, or blocking their downstream signaling, after acute infection could provide therapeutic benefit in PASC. Additionally, PASC may be driven by activation of stress response programs as evidenced by upregulated AP-1 and *DDIT4* with elevated IL-1β signaling, which could result from an imbalance between autophagy and inflammasome activation. Overall, network analyses provide a platform for generating mechanistic hypotheses on key pathways in mild COVID-19 from acute infection through resolution that may prime adaptive immune responses, mediate successful resolution of infection, and enable diagnosis and treatment of complications such as PASC.

## Discussion

Our study provided an in-depth longitudinal analysis of the immune response to SARS-CoV-2 natural infection and mild COVID-19 by integrating serum proteomics, single-cell transcriptomics and epigenomics, and cellular immunophenotype by flow cytometry with clinical metadata and comprehensive analysis of the SARS-CoV-2-specific adaptive immune response in T cells, memory B cells, and antibodies. To our knowledge, this is the deepest longitudinal systems immunology study to-date in mild COVID-19, and reveals numerous new insights. First, we define the immune response to early acute infection including inflammatory cytokine and innate sensor signaling, stronger IFN responses in older participants, and a potential IFN-plasmablast regulatory circuit. A subset of these changes were correlated with the humoral response to SARS-CoV-2 in convalescence. We then confirmed that the longitudinal resolution of these inflammatory pathways was coordinated with re-establishment of homeostasis in most participants. Five PASC participants were exceptions to this resolution, and could be distinguished by dampened IFN and antiviral responses during acute infection coupled with prolonged inflammation. Finally, we integrated these data to identify potential master regulatory nodes in early infection, longitudinal resolution, and PASC for hypothesis generation and validation of biomarkers and therapeutics.

A defining characteristic in our mild COVID-19 cohort was robust immune activation in the first 2 weeks of acute infection that resolved over time. This included inflammatory cytokine responses (IFNs, TNF), innate immune sensor signaling pathways, and activation in adaptive and innate immune cells. The key innate immune sensors triggered in natural SARS-CoV-2 infection are not confirmed, but multiple data types strongly implicate RIG-I. Serum proteomics identified increased extracellular levels of multiple PRR pathway members including RIG-I during acute infection, but their source and function are unclear. These may derive from recent cell death or extracellular vesicles, which have been reported to potentially transfer TLRs between cells (Zhang et al., 2019). Innate danger sensors are key drivers of the IFN response, which was also robustly induced in our cohort along with other inflammatory signals such as TNF. A subset of upregulated proteins (SAMD9L, CXCL10, CXCL11) is selective to type I/II IFN responses, suggesting bias away from type III IFNs systemically (Allenspach et al., 2021; Forero et al., 2019). As these pathways waned over time, activation marker-positive cells and inflammatory proteins largely returned to uninfected control levels around day 30 PSO. This temporal control is likely critical for successful resolution of mild COVID-19. This contrasts with persistent CRS reported in severe COVID-19, which includes mechanisms of inflammatory damage to tissue such as TNF/IFNγ-mediated cell death (Karki et al., 2021). Proteins involved in homeostatic functions (EMT, coagulation, angiogenesis) increased from acute infection to convalescence. The longitudinal increase of multiple coagulation pathway proteins may contribute to reported increases in risk of immune thrombocytopenia, a complication associated with severe COVID-19 infection (Guan et al., 2020). We found levels of THBD were significantly increased in convalescence of mild COVID-19, and has been reported to strongly correlate with duration of hospitalization and risk of mortality in hospitalized COVID-19 (Goshua et al., 2020). These results suggest a link between the inflammatory response in acute infection, the kinetics of inflammatory resolution, and their dysregulation in long-term coagulopathy risk and severe COVID-19.

We observed a clear increase in immune responses with advancing age in mild COVID-19. Age is among the strongest risk factors for severe COVID-19 and mortality, but the mechanisms underlying these effects remain poorly defined (Williamson et al., 2020). Many studies are confounded by age when comparing uninfected controls and mild COVID-19 cases, which are often younger, with typically older individuals in moderate and severe COVID-19. A male-specific age effect was correlated to poorer CD8+ T cell responses and disease severity with advanced age (Takahashi et al., 2020). Our cohort was age-matched between COVID-19 participants and uninfected controls, and age was not significantly correlated with illness severity score. However, IFN responses showed a dramatic age-related effect: increased responsiveness of PBMCs from older COVID-19 participants to IFNs, as evidenced by both more pathway enrichment and higher enrichment scores from scRNAseq, and cell type-specific enrichment of IRF motifs in scATACseq. Older participants also showed increased cytokine and danger sensor signaling in innate immune cells. Enhanced IFN responses and TLR signaling in older participants are contrary to expectations from prior studies, where advanced age is associated with impaired IFN and TLR responses and age-associated decrease of TLR responses in DCs (Molony et al., 2017; Panda et al., 2010; Pillai et al., 2016). These observations could be due to higher viral loads in older participants (Euser et al., 2021; Mahallawi et al., 2021), cells from older participants being more intrinsically reactive to cytokines, the magnitude of inflammatory cytokine response being higher, or persistence of inflammatory cytokines longer in older participants. We also observed increased adaptive immune cell activation, which contrasts with expectations from prior studies showing immunosenescence in elderly healthy individuals. Differences between our results and prior studies on age-related effects in immunity may also be due to our cohort including more young and middle-aged adults <55 years old compared to elderly adults typically >65 years old. Collectively, our results indicate that SARS-CoV-2 infection triggers enhanced inflammatory responses in older individuals in mild COVID-19, which may underlie the increased risk for severe COVID-19 in older populations.

Robust plasmablast expansion is a feature of viral infections (dengue, Ebola), vaccines, and chronic autoimmune diseases (Kim et al., 2016; McElroy et al., 2015; Wrammert et al., 2012). Previous studies in moderate and severe COVID-19 reported robust plasmablast and extrafollicular B cell responses (Bernardes et al., 2020; Kaneko et al., 2020; Kuri-Cervantes et al., 2020; Mathew et al., 2020; Ren et al., 2021; Stephenson et al., 2021; Woodruff et al., 2020). We also observed a robust plasmablast expansion in acute infection. A fraction of these plasmablasts were CoV-2 spike protein-specific, indicating these cells can in principle contribute to early control of viral replication and are at least partly CoV-2-specific. PD-1^high^ CXCR5-Tph cells were also positively correlated with plasmablasts, potentially providing help as observed in autoimmune diseases (Rao et al., 2017). IFN responses are among the key drivers of extrafollicular B cell subsets in autoimmunity (Manni et al., 2018; Soni et al., 2020), which was consistent with our scRNAseq data showing enhanced IFN signaling and our scATACseq data showing IRF motif enrichments in plasmablasts. The functional consequence of these plasmablasts in SARS-CoV-2 infection remains unclear, but may connect early IFN responses to antibody titers.

Priming of adaptive immunity is critical to successful resolution of acute infection and protection against reinfection. Correlates from acute infection were identified that explain interindividual heterogeneity in magnitude of humoral immune responses. Serum proteins involved in innate immune pathways, including IFN responses, chemokines, and PRR signaling, were positively correlated with RBD IgG titer, neutralizing antibody titer, and CoV-2-specific memory B cell frequency. These were also positively correlated with activation of T and B cells, indicating the importance of coordination in acute infection for an optimal humoral response. Complement proteins were also enriched, consistent with prior studies showing complement can facilitate antigen retention in follicular DCs (Phan et al., 2007) and enhancing BCR-mediated signaling (Fischer et al., 1998; Lyubchenko et al., 2005). Plasmablasts were a positive correlate of antibodies and memory B cells, but it is unclear whether they play a functional role in clearing infection. Overall, these findings demonstrate that the immune response to acute infection is critical to an effective humoral response, emphasizing the importance of coordination between innate and adaptive arms of immunity.

PASC or long COVID is one of the most enigmatic consequences of the ongoing pandemic. The involvement of many organ systems coupled with the highly subjective nature of symptoms has made it difficult to define objective consensus criteria for diagnosis or clear therapeutic options. In our multi-omics discovery cohort, a subset of five COVID-19 participants progressed to PASC. All PASC participants in this discovery cohort were female, consistent with prior reports of female-biased presentation. Previous studies have suggested that females may have stronger inflammatory responses to vaccines and infections, and predisposition to autoimmune disease (Klein and Flanagan, 2016)) but it is unclear whether this could contribute to the skewed incidence of PASC in females. Significant correlation was observed between PASC and number of initial symptoms as well as illness severity score during acute infection, as previously reported (Blomberg et al., 2021; Sudre et al., 2021). Select SARS-CoV-2-specific humoral responses were negatively correlated with PASC, but previous studies report both positive and negative associations of SARS-CoV-2-specific antibody titers and T cell responses with PASC (Blomberg et al., 2021; Files et al., 2021; García-Abellán et al., 2021). Larger studies are likely required to clarify these correlations.

Our expanded PASC cohort was more diverse (60 female, 26 male) and validated the inflammatory protein signatures identified in serum that distinguished PASC participants from recovered COVID-19 participants after acute infection, particularly in those with the most severe symptoms. These signatures were coupled with evidence of persistent activation and inflammatory cytokine signaling in innate immune cells based on single-cell gene expression and chromatin accessibility after 30 days PSO. DCs and CD14+ monocytes in PASC showed the most differences in transcription factor motif accessibility enrichment. Among these, AP-1, STAT, IRF, BATF, and BACH suggest ongoing cellular stress, immune cell activation and differentiation, and inflammatory cytokine signaling during PASC. Gene expression signatures demonstrated increases in immunologically important transcripts from genes proximal to AP-1 sites that were differentially accessible in PASC monocytes. Furthermore, transcriptomics showed stronger TNF signaling in CD14+ monocytes from PASC participants. Early infection signaling and kinetics were also unique in PASC, including lower IFN responses in acute infection that did not wane longitudinally. This combination of changes mirrors severe COVID-19: dampened antiviral responses may fail to control viral replication in both, which can drive innate immune responses to persist beyond acute infection and cause ongoing pathology.

Integrative analysis defined a PASC-specific inflammatory cytokine signaling landscape that suggests multiple therapeutic targets. Single-cell gene expression and serum proteome data provided evidence of persistently elevated IFNγ, IFNλ1, IL-1β, IL-12, IL-6, and TNF in PASC. scATACseq provided evidence of active signaling from these cytokines, with increased accessibility of motifs for key transcription factors in PASC, including IRFs, STATs, and AP-1 in innate immune cell types, primarily DCs and CD14+ monocytes. Cumulatively, these findings suggest blockade of persistent inflammatory cytokine signaling may be therapeutic in PASC. JAK inhibitors are an appealing therapeutic option due to their ability to dampen signaling from several of the cytokines found to be elevated in PASC in addition to their short half-life that facilitates the ability to perform short therapeutic trials to assess efficacy. Similarly, the short half-life and safety of the IL-1 receptor antagonist anakinra could be a strong candidate for early therapeutic trials to probe the role of IL-1 in driving disease. Both JAK inhibitors and IL-1 receptor blockade have been tested in the context of severe COVID-19 immunopathology, with mixed results. Persistently elevated TNF may also be an appealing target given the potential for TNF-driven pathogenic cell death and correlations with COVID-19 disease severity and mortality (Del Valle et al., 2020; Karki et al., 2021). TNF can also drive IL-1β expression, along with AP-1 and STAT/IRF enrichment observed in motif analyses, motivating therapeutic inhibition of TNF in PASC through TNF blockade. Another promising target is IFNγ, which was significantly elevated in serum from PASC participants compared to recovered participants, coexpressed among pathways in CD14+ monocytes by scRNAseq, and was a strongly predicted ligand signal from ligand-receptor interactions in the majority of cell types. Both TNF and IFNγ are also associated with severe COVID-19, suggesting similarities between severe disease and PASC. Broader anti-inflammatory therapy such as corticosteroids may be useful to attenuate pathogenic inflammation in PASC and has been tested in COVID-19 patients with persistent inflammatory lung disease (Myall et al., 2021) but chronic steroid use is associated with many side effects that make effective targeted therapies more desirable. These novel insights into PASC can focus future studies on pathogenic mechanisms and evaluating efficacy of therapeutic targets. Molecular classification of disease heterogeneity in PASC may identify disease subsets to further enhance clinical management and optimize therapeutic strategies.

There are several key limitations to our study, including 1) small sample size, especially for sickest PASC, 2) lack of geographic and racial diversity, 3) asynchronous sampling due to outpatient status, and 4) gaps in sampling during early infection (first ~7 days PSO) for some participants. Natural history studies are intrinsically limited to correlative associations. Confirming causality from our findings to mechanistically link early infection immune responses to convalescent CoV-2-specific immune responses will require preclinical models. Multi-omic assays were conducted independently from CoV-2-specific assays on parallel samples from each blood draw, so relationships between different -omics and CoV-2 responses were inferred. Direct multiplexed analysis, particularly adding TCR and BCR clonality, will better enable dissection of virus-specific vs. non-specific immune mechanisms. Despite these weaknesses, our study provides insightful data that advance our understanding of SARS-CoV-2 infection, mild COVID-19, convalescence, and PASC.

Overall, our study results provide a comprehensive longitudinal roadmap of immune activation and resolution in mild COVID-19, including a potential mechanism for an age-dependent effect on immune responses. We observed a robust plasmablast response that may be tightly regulated by early IFN responses, and identified key early correlates of antibody and B cell responses, both findings which should be broadly tested as potential shared features in diverse natural infections. A subset of participants who progressed to PASC revealed novel inflammatory and non-inflammatory signatures in serum proteins, and innate immune-centric hyperactivation. Multiple potential therapeutic targets in PASC are nominated by our analyses, and serum protein biomarkers may provide opportunities for objective diagnosis of inflammatory and non-inflammatory PASC after validation in larger cohorts. A more personalized approach to immunomonitoring and therapy has the potential to improve outcomes across the spectrum of COVID-19 and PASC.

## Acknowledgements

We thank the study participants for their dedication to this project; the Allen Institute founder, Paul G. Allen, for his vision, encouragement, and support; Ken Stuart, for constructive feedback on early drafts; the Human Immune System Explorer (HISE) software development team at the Allen Institute for Immunology for their support and dedication. This paper and the research behind it would not have been possible without the collaborative computational data analysis environment provided by HISE.

## Supplemental figure legends

**Fig. S1: A)** Principal component analysis (PCA) bi-plot of the serum proteome across all longitudinal COVID-19 participant samples (red points) and uninfected controls (green points). The loadings of the most varying proteins are represented by arrows; arrow length is proportional to the variance of the proteins contributing to sample distinction. Shapes of points represent sex and age represents age. A Wilcoxon rank-sum test was used to test if the distribution of principal components 1 and 2 was significantly different between COVID-19 participants and uninfected controls (p-value < 0.05). **B)** PCA of flow cytometry cell frequencies across all longitudinal COVID-19 participant samples and uninfected controls, including batch bridging controls. Points are colored by batch ID. Principal Variance Component Analysis (PVCA) was performed as indicated in the bar plot to assess batch effects. The proportion of variance (y-axis) in cell frequencies contributed by each factor (x-axis) is depicted. **C)** PCA bi-plot of the flow cytometry-based proportions across all longitudinal COVID-19 participant samples (red points) and uninfected controls (green points). The loadings of the top varying cell proportions are represented by arrows. A Wilcoxon rank-sum test was used to test if principal components 1 and 2 significantly (p-value < 0.05) differentiate infection status. **D)** UMAP density representation per batch as grids and PVCA analysis (bar plot) of the scRNAseq batch bridging control samples. The proportion of variance (y-axis) in gene expression contributed by each factor (x-axis) is depicted in the bar plot. **E)** UMAP density representation per batch as grids and PVCA analysis (bar plot) of the full scRNAseq cohort samples including all COVID-19 longitudinal samples and uninfected controls. The proportion of variance (y-axis) in gene expression contributed by each factor (x-axis) is depicted in the bar plot. **F)** UMAP of the cell type annotation on scRNAseq data using the Weighted Nearest Neighbors (WNN) approach implemented in Seurat v4 with a CITE-seq based reference dataset (Hao et al., 2020b). Each point represents single cells color-coded by cell type label. **G)** Heatmaps of selected gene and predicted ADT (x-axes) expression by cell type (y-axes) of scRNAseq data. **H)** Correlation scatter plots between scRNA proportions (x-axis) and the equivalent population gating in flow cytometry (y-axis) across all samples. Significance of correlations was tested by a Spearman correlation test and only significant correlations are shown.

**Fig. S2: A)** Box and jitter plots of scRNAseq-based cell type proportions that were significantly different (adjusted p-values < 0.05) comparing the early acute infection timepoints of COVID-19 participants (red) to uninfected controls (green) assessed by fitting a linear model adjusted for age and sex. **B)** Heatmap of the flow cytometry and scRNAseq cell type proportions (rows) that were significantly different (adjusted p-values < 0.05 assessed by Wilcoxon rank-sum test) comparing older (including middle aged and senior adults) to younger COVID-19 participants (columns) in early acute infection. **C)** Heatmap of serum proteins (rows) that were significantly different (adjusted p-values < 0.05 assessed by Wilcoxon rank-sum test) comparing older to younger COVID-19 participants (columns) in early acute infection. **D)** Bar plot of the number of DEGs (y-axis) comparing age-matched early acute COVID-19 participants to uninfected controls (per grid) within each cell type (x-axis) detected in scRNAseq. Number of DEGs that were significantly up-regulated (red) and down-regulated (blue) at an adjusted p-value < 0.05 are reported. **E)** GSEA pathway enrichment (y-axis) among DEGs comparing age matched early acute COVID-19 participants to uninfected controls (per grid) in every cell type (x-axis) of scRNAseq data. The color gradient of points represents the NES per pathway indicating up-regulation (red) or down-regulation (blue) of each pathway. The size of points represents the percent of genes enriched in a pathway per cell type and age group. **F)** GSEA pathway enrichment (y-axis) among DEGs comparing early acute infection timepoints of older COVID-19 participants to younger in every cell type (x-axis) of scRNAseq data. The color gradient of points represents the positive NES per pathway indicating up-regulation of the pathways in older participants. The size of points represent the percent of genes enriched in a pathway per cell type and age group. Pathways are arranged as being expressed in most to the least number of cell types. **G)** Scatterplot of the -log10 of the lowest P value per cell type by the absolute value of the maximum z-score shift per cell type. CD16+ monocytes, CD14+ monocytes, DCs, and plasmablasts appear to be the most impacted during the early infection period. **H)** Heatmap of top 50 motif accessibility changes during early COVID19 infection. Z-scores representing motif accessibility as compared to background were calculated for each cell. Z-score shifts were tested for statistical significance by a Wilcoxon rank-sum test with an FDR < 0.1. Only significant shifts are colored.

**Fig. S3: A)**. Serum cytokines, chemokines and complement proteins (individual grids) in COVID-19 participants that significantly (adjusted p-values < 0.05) decreased over time. Individual lines are color-coded by the participant ID. **B)** scRNA based cell type proportions (individual grids) among COVID-19 participants that significantly (adjusted p-values < 0.05) showed decrease over time. Individual lines are color coded by the participant ID. **C)** Pairwise correlations between functional pathways (circles, serum proteome) and cell type proportions (squares, flow cytometry; triangles, scRNAseq) overtime. The edge color gradient indicates significant (p-values < 0.05) positive (red) or negative (blue) Spearman correlations over time. **D)** GSEA pathway enrichment (y-axis) among longitudinal gene changes per COVID-19 participant in every cell type (x-axis) of scRNAseq data. The size of points represents the percent of COVID-19 participants the pathway is enriched in. Pathways are arranged as being expressed in most to the least number of cell types across participants. **E)** Longitudinal IRF motif accessibility per participant over disease course. Curves are colored for PASC and sickest PASC (orange and red respectively) and recovered (black) COVID-19 participants. Loess curves were added to visualize group trends. Uninfected participants are represented by black solid circles on the left. **F)** PCoA of serum proteome trajectories for individual COVID-19 participants and uninfected controls (black circles, no arrows). Symbols are colored by comorbidities and clinical features. Arrows are colored by visit interval over time.

**Fig. S4: A)** Bar plot of the number of DEGs (y-axis) comparing early acute infection timepoints of PASC participants to recovered COVID-19 participants in every cell type (x-axis) detected in scRNAseq. Number of DEGs that were significantly up-regulated (red) and down-regulated (blue) at an adjusted p-value < 0.05 are reported. **B)** Box plot of the SLEA score (y-axis) of the interferon response pathways in scRNAseq significantly downregulated (adjusted p-values < 0.05) in CD8+ TEMRAs and TEM from PASC compared to recovered COVID-19 participants in early acute infection. The SLEA score of the pathways per sample was calculated and tracked over the course of disease (x-axis). **C)** Arc co-expression network of genes enriched in the interferon responses pathway in CD14+ monocytes, from early acute timepoints of recovered participants. Edges represent correlation between genes at greater than 0.8 Spearman correlation coefficient, while size of each gene node represents the degree or number of gene correlations above mentioned coefficient. **D)** Arc co-expression network of genes enriched in the IL1B and TLR signaling pathways in CD14+ monocytes, from early acute timepoints of PASC participants. Edges represent correlation between genes at greater than 0.8 Spearman correlation coefficient, while size of each gene node represents the degree or number of gene correlations above mentioned coefficient. **E)** Bar plot of the pathways (y-axis) enriched in CD14+ monocytes comparing PASC to recovered participants in early acute infection (red bars) and beyond 30 days PSO (black bars). X-axis represents the GSEA NES, with a positive NES indicating up-regulation in PASC, while negative NES indicating down-regulation in PASC compared to recovered. **F)** Box plot of the SLEA score (y-axis) of TNF signaling via NFKb pathway significantly upregulated (adjusted p-values < 0.05) in CD8+ TEM and TEMRAs from PASC compared to recovered COVID-19 participants in early acute infection. The SLEA score of the pathways per sample was calculated based on scRNAseq gene expression per cell type and tracked over the course of disease (x-axis). **G)** Number of differentially expressed serum proteins (y-axis) over time (x-axis) compared to uninfected controls for each COVID-19 participant. Outlier analysis was performed comparing COVID-19 participants to uninfected background and selecting features >2 standard deviations from the mean. Recovered, PASC and the sickest PASC COVID-19 participants are colored by grey, orange and red respectively. **H)** Heatmap of the Gene Set Enrichment Analysis results in scRNAseq beyond 30 days PSO among a curated AP-1 target gene set. These genes were identified as those nearest to differentially accessible scATAC-seq peaks with AP-1 motifs. Columns represent pathways while rows represent the genes enriched among pathways. The inset on the right shows genes unique to the “innate immune system” pathway. **I)** Ligand activity observed in CD14+ monocytes and their expression in sender cell types of PASC participants, **J)** dot plot showing expression of these ligands among PBMCs cell types, where the size of circles indicates the percentage of cells expressing the gene for each ligand and the color scale indicates the average expression level, and **K)** target genes of predicted ligands to CD14+ monocytes.

**Fig. S5. A)** Correlation scatterplots between the observed (y-axis) and estimated values (x-axis) as obtained by fitting linear mixed effect models over time to the serum proteome and **B)** to flow cytometry based proportions. For flow cytometry, the top 15 (ranked by Spearman’s correlation coefficient) are shown. All correlations were significant (p-values < 0.05) with greater than a 0.5 coefficient as assessed by the Spearman’s correlation test. **C)** Day 7 estimated serum proteins and pathway enrichments, and cell proportions **(D)** that significantly correlated (Spearman’s correlation coefficients > 0.5 and adjusted p-values < 0.05) with the estimated S-specific memory B cells. Rows correspond to proteins or cell frequencies measured by flow cytometry. Columns correspond to individual COVID-19 participants arranged by increasing order outcomes (represented as column annotation bars).

**Fig. S6**. Differentially expressed immune-related proteins observed in early acute infection (≤15 days PSO), longitudinally and in PASC COVID-19 participants from serum proteomics. All samples from the broader serum proteomics cohort are represented in the heatmap.

**Fig. S7. A)** Gene expression of differentially expressed serum proteins per cell type. The scaled average expression shown for each gene in a given cell type in COVID19 participants. The size of the circle represents the percent cells expressing genes. **B)** Serum protein expression changes in COVID-19 participants (recovered and PASC) are shown in the box plot. Samples were grouped based on days PSO covering: 1) early acute infection ≤15 days; 2) late acute infection, 16-30 days; and 3) post-acute infection and convalescence, ≥31 days. Wilcoxon rank-sum tests were used to compare the PASC to recovered participants across all timepoints, and a representative subset of significantly differential proteins are shown. P-values < 0.05 were considered statistically significant.

## METHODS

### RESOURCE AVAILABILITY

#### Lead contacts

Further information and requests for resources and reagents should be directed to and will be fulfilled by the lead contacts: Gregory Szeto; M. Juliana McElrath; and Thomas F. Bumol.

#### Materials Availability

This study did not generate new unique reagents.

#### Data and code availability

The RNAseq and ATACseq data generated during this study are available at GEO under accession number GSE173590. All analytical code and figure generation will be available on Github.

### EXPERIMENTAL MODEL AND SUBJECT DETAILS

#### Demographics

Eighteen immunocompetent study participants who were diagnosed with SARS-CoV-2 were selected for these studies, as they had donated at least one sample of serum at or earlier than 15 days post-symptom onset. Both studies were recruited at the Seattle Vaccine Trials Unit (Seattle, Washington, USA). The cohort demographics are described in the following table.

| Volunteers with SARS CoV-2 Diagnosis | Females | Males |
| --- | --- | --- |
| Number | 10 | 8 |
| Age | Mean 46<br>Range 29-79 | Mean 46<br>Range 22-65 |
| Race and Ethnicity | 100% White (n=10) | 100% White (n=8)<br>12.5% Hispanic/Latino (n=1) |
| Co-morbidities previous to SARS-CoV2 Diagnosis | Asthma (n=1)<br>High Blood Pressure (n=1)<br>Autoimmune Disease (n=2)<br>Current or past Smoker (n=1)<br>Anemia (n=1)<br>Obesity (n=2) | Heart disease (n=1)<br>High Blood Pressure (n=1)<br>Current or past Smoker (n=1)<br>Obesity (n=1) |

| Uninfected SARS-CoV-2-negative Volunteers | Females | Males |
| --- | --- | --- |
| Number | 10 | 13 |
| Age | Mean 47<br>Range 29-61 | Mean 48<br>Range 31-77 |
| Race and Ethnicity | 100% White (n=10) | 84.6% White (n=11)<br>15.4% Hispanic/Latino (n=2)<br>Other:<br>American Indian or Alaska Native;White (n=1)<br>Asian;White (n=1) |
| Co-morbidities previous to SARS-CoV2 Diagnosis | Asthma (n=1)<br>High Blood Pressure (n=1)<br>Autoimmune Disease (n=3)<br>Current or past Smoker (n=3)<br>Anemia (n=1) | COPD (n=1)<br>Cancer (n=1)<br>Current or past Smoker (n=2)<br>High Blood Pressure (n=1)<br>Hypertension(n=1) |

Informed consent was obtained from all participants and the Fred Hutchinson Cancer Research Center Institutional Review Board approved the study and procedures (IR10440).

### METHODS DETAILS

#### Study Conduct

Peripheral blood mononuclear cells (PBMCs) and serum were collected from participants enrolled in the longitudinal study, “Seattle COVID-19 Cohort Study to Evaluate Immune Responses in Persons at Risk and with SARS-CoV-2 Infection”. Eligibility criteria included adults in the greater Seattle area at risk for SARS-CoV2 infection or those diagnosed with COVID-19 by a commercially available SARS CoV-2 PCR assay. Study data were collected and managed using REDCap electronic data capture tools hosted at Fred Hutchinson Cancer Research Center, including detailed information on symptoms during acute infection and longitudinal follow-up ranging from 33-233 days post symptom onset. Plasma from pre-pandemic controls used for ELISA controls were blindly selected at random from the study, “Establishing Immunologic Assays for Determining HIV-1 Prevention and Control”, with no considerations made for age, or sex. Informed consent was obtained from all participants at the Seattle Vaccine Trials Unit and the Fred Hutchinson Cancer Research Center Institutional Review Board approved the studies and procedures.

#### Regulatory approvals from FH and AIFI

##### COVID19 FH samples and healthy controls

FH RG: 1007696 IR File: 10440 Main Consent 04/05/2020 and 6/04/2020 Seattle COVID-19 Cohort Study to Evaluate Immune Responses in Persons at Risk and with SARS-CoV-2 Infection.

##### Batch control leukopack from BIOIVT used in flow cytometry

PROSPECTIVE COLLECTION OF NON-MOBILIZED LEUKOCYTES VIA LEUKAPHERESIS FOR RESEARCH; PROTOCOL NO: SERATRIALS-18002; WIRB® Protocol #20190318

#### Serum and PBMC isolation

Blood collected in SST tubes was allowed to clot for at least 30 min at room temperature. Following cotting, the samples were centrifuged in the primary tubes at 1600-1800 X for 10 min. Serum was transferred to a 15ml tube, aliquoted into 300ul aliquots and stored at −80°C until use in the assays.

Blood collected in acid citrate dextrose tubes was transferred to Leucosep tubes (Greiner Bio One). The tube was centrifuged at 800-1000g for 15 min and the PBMC layer recovered above the frit. Peripheral blood mononuclear cells (PBMCs) were washed twice with Hanks Balanced Solution without Ca+ or Mg+ (Gibco) at 200-400xg for 10 min, counted, and aliquoted in heat-inactivated fetal bovine serum with 10% dimethylsulfoxide (DMSO, Sigma) for cryopreservation. PBMCs were cryopreserved at −80°C in Stratacooler (Nalgene) and transferred to liquid nitrogen for long-term storage.

#### Cell Type Flow Cytometry

To assess cell type proportions, PBMCs were analyzed with four 25-color immunophenotyping flow cytometry panels (P1, P2, P3 and P4). 1×10^6^ thawed PBMCs were centrifuged (750×g for 5 minutes at 4°C) using a swinging bucket rotor (Beckman Coulter Avanti J-15RIVD with JS4.750 swinging bucket, B99516), the supernatant was removed using a vacuum aspirator pipette, and the cell pellet resuspended in DPBS without calcium and magnesium (Corning 21-031-CM). Cells were incubated with Fixable Viability Stain 510 (BD, 564406) and an Fc receptor blocking reagent, TruStain FcX (BioLegend, 422302) P1, P3 and P4 or purified mouse IgG (Bio-Rad, PMP01) P2 for 30 minutes at 4°C, then washed with chilled Cell Staining Buffer (BioLegend, 420201). Cells were stained with a cocktail of antibodies (P1, P2, P3 and P4) in Cell Staining Buffer plus BD Horizon Brilliant Stain Buffer Plus (BD, 566385) (**Key Resources Table**) at a staining volume of 100 μl for 30 minutes at 4°C, then washed with chilled Cell Staining Buffer. Fixation was performed by resuspending cells in 100 μl of FluoroFix Buffer (BioLegend, 422101) and incubating for 30 minutes at 25°C, protected from light. Following fixation, cells were washed twice with Cell Staining Buffer and resuspended in 100 μl Cell Staining Buffer. Stained cells were analyzed on a 5 laser Cytek Aurora spectral flow cytometer. Spectral unmixing was calculated with pre-recorded reference controls using Cytek SpectroFlo software (Version 2.0.2). Cell types were quantified by traditional bivariate gating analysis performed with FlowJo cytometry software (Version 10.7)

#### Spike and RBD Memory B cell flow cytometry assays

Fluorescent SARS-CoV-2-specific S6P (Hsieh et al., 2020) (provided by Roland Strong, Fred Hutchinson Cancer Research Center, Seattle, WA) and RBD (provided by Leonidas Stamatatos, Fred Hutchinson Cancer Research Center, Seattle, WA) probes were made by combining biotinylated protein with fluorescently labeled streptavidin (SA). The S6P probes were made at a ratio of 1:1 molar ratio of trimer to SA. Two S6P probes, one labeled with AlexaFluor488 (Invitrogen), one labeled with AlexaFluor647 (Invitrogen), were used in this panel in order to increase specificity of the detection of SARS-CoV-2-specific B cells. The RBD probe was prepared at a 4:1 molar ratio of RBD monomers to SA, labeled with R-phycoerythrin (Invitrogen). Cryopreserved PBMCs from SARS-CoV-2-convalescent participants and a pre-pandemic SARS-CoV-2-naïve participants were thawed at 37°C and stained for SARS-CoV-2-specific memory B cells as described previously (Seydoux et al., 2020) with a flow cytometry panel shown in Reagents Table 1. Cells were stained first with the viability stain (Invitrogen) in PBS for 15 min at 4°C. Cells were then washed with 2% FBS/PBS and stained with a cocktail of the three probes for 30 min at 4°C. The probe cocktail was washed off with 2% FBS/PBS and the samples were stained with the remaining antibody panel and incubated for 25 min at 4°C. The cells were washed two times and resuspended in 1% paraformaldehyde/1x PBS for collection on a LSR II or FACSymphony flow cytometer (BD Biosciences). Data was analyzed in Flow Jo version 9.9.4.

#### Intracellular Cytokine Staining (ICS) Assay

Flow cytometry was used to examine SARS-CoV-2-specific CD4+ and CD8+ T-cell responses using a validated ICS assay. The assay was similar to a published report (Dintwe et al., 2019; Horton et al., 2007) and the details of the staining panel are included in Reagents Table 2. Peptide pools covering the structural proteins of SARS-CoV-2 were used for the six-hour stimulation. Peptides matching the SARS-CoV-2 spike sequence (316 peptides, plus 4 peptides covering the G614 variant) were synthesized as 15 amino acids long with 11 amino acids overlap and pooled in 2 pools (S1 and S2) for testing (BioSynthesis). All other peptides were 13 amino acids overlapping by 11 amino acids and were synthesized by GenScript. The peptides covering the envelope (E), membrane (M) and nucleocapsid (N) were initially combined into one peptide pool, but the majority of the assays were performed using a separate pool for N and one that combined only E and M. Several of the open reading frame (ORF) peptides were combined into two pools, ORF 3a and 6, and ORF 7a, 7b and 8. All peptide pools were used at a final concentration of 1 microgram/ml for each peptide. As a negative control, cells were not stimulated, only the peptide diluent (DMSO) was included. As a positive control, cells were stimulated with a polyclonal stimulant, staphylococcal enterotoxin B (SEB). Cells expressing IFNγ and/or IL-2 and/or CD154 were the primary immunogenicity endpoint for CD4+ T cells and cells expressing IFNγ were the primary immunogenicity endpoint for CD8+ T cells. The overall response to SARS-CoV-2 was defined as the sum of the background-subtracted responses to each of the individual pools. A sample was considered positive for CD4+ or CD8+ T cell responses to SARS-CoV-2 if any of the CD4+ or CD8+ T cell responses to the individual peptide pool stimulations was positive. Positive responses to a given peptide pool stimulation were determined using the MIMOSA (Mixture Models for Single-Cell Assays) method (Finak et al., 2014a). The MIMOSA method uses Bayesian hierarchical mixture models that incorporate information on cell count and cell proportion to define a positive response by comparing peptide-stimulated cells and unstimulated negative controls. MIMOSA estimates the probabilities that peptide-stimulated responses are responders and applies a false-discovery rate multiplicity adjustment procedure (Newton et al 2004). Responses with false-discovery rate q-values < 0.05 were considered positive.The total number of CD4+ T cells must have exceeded 10,000 and the total number of CD8+ T cells must have exceeded 5,000 for the assay data to be included in the analysis.

#### Antibody ELISAs for RBD and N

Half-well area plates (Greiner) were coated with purified RBD protein at 16.25ng/well in PBS (Gibco) for 14-24h at room temperature. After 4 150ul washes with 1X PBS, 0.02% Tween-2 (Sigma) using the BioTek ELx405 plate washer, the IgA and IgG plates were blocked at 37°C for 1-2 hours with 1X PBS, 10% non-fat milk (Lab Scientific), 0.02% Tween-20 (Sigma); IgM plates were blocked with 1X PBS, 10% non-fat milk, 0.05% Tween-20.

Serum samples were heat inactivated by incubating at 56°C for 30 minutes, then centrifuged at 10,000 x g / 5 minutes, and stored at 4°C previous to use in the assay. For IgG ELISAs, serum was diluted into blocking buffer in 7-12 1:4 serial dilutions starting at 1:50. For IgM and IgA ELISAs, serum was diluted into 7 1:4 serial dilutions starting at 1:12.5 to account for their lower concentration. A qualified pre-pandemic sample (negative control) and a standardized mix of seropositive serums (positive control) was run in each plate and using to define passing criteria for each plate. All controls and test serums at multiple dilutions were plated in duplicate and incubated at 37°C for 1 hour, followed by 4 washes in the automated washer. 8 wells in each plate did not receive any serum and served as blocking controls.

Plates then were plated with secondary antibodies (all from Jackson ImmunoResearch) diluted in blocking buffer for 1h at 37C. IgG plates used donkey anti-human IgG HRP diluted at 1:7500; IgM plates used goat anti-human IgM HRP diluted at 1:10,000; IgA plates used goat anti-human IgA HRP at 1:5000. After 4 washes, plates were developed with 25ul of SureBlock Reserve TMB Microwell Peroxide Substrate (Seracare) for 4 min, and the reaction stopped by the addition of 50ml 1N sulfuric acid (Fisher) to all wells. Plates were read at OD_450nm_ on SpectraMax i3X ELISA plate reader within 20 min of adding the stop solution.

OD_450nm_ measurements for each dilution of each sample were used to extrapolate RBD endpoint titers when CVs were less than 20%. Using Excel, endpoint titers were determined by calculating the point in the curve at which the dilution of the sample surpassed that of 5 times the average OD_450nm_ of blocking controls + 1 standard deviation of blocking controls.

N IgG was measured using SARS CoV-2 IgG Architect (Abbot), and index values used as a quantitative measure of the N binding activity.

#### Neutralizing antibody assay

##### Viruses and cells

VeroE6 cells were obtained from ATCC (clone E6, ATCC, #CRL-1586) and cultured in complete DMEM medium consisting of 1x DMEM (VWR, #45000-304), 10% FBS, 25mM HEPES Buffer (Corning Cellgro), 2mM L-glutamine, 1mM sodium pyruvate, 1x Non-essential Amino Acids, and 1x antibiotics. The infectious clone SARS-CoV-2 (icSARS-CoV-2-mNG), derived from the 2019-nCoV/USA_WA1/2020 strain, was propagated in VeroE6 cells (ATCC) and sequenced (Xie et al., 2020).

##### Focus Reduction Neutralization Test

Neutralization assays with SARS-CoV-2 virus were performed as previously described (Vanderheiden et al., 2020b). Plasma/serum were serially diluted (three-fold) in serum-free Dulbecco’s modified Eagle’s medium (DMEM) in duplicate wells and incubated with 100–200 FFU infectious clone derived SARS-CoV-2-mNG virus at 37°C for 1 hr. The antibody-virus mixture was added to VeroE6 cell (C1008, ATCC, #CRL-1586) monolayers seeded in 96-well blackout plates and incubated at 37°C for 1 h. Post-incubation, the inoculum was removed and replaced with pre-warmed complete DMEM containing 0.85% methylcellulose. Plates were incubated at 37°C for 24 h. After 24 h, methylcellulose overlay was removed, cells were washed twice with PBS and fixed with 2% paraformaldehyde in PBS for 30 min at room temperature. Following fixation, plates were washed twice with PBS and foci were visualized on a fluorescence ELISPOT reader (CTL ImmunoSpot S6 Universal Analyzer) and enumerated using Viridot (Katzelnick et al., 2018). The neutralization titers were calculated as follows: 1 - (ratio of the mean number of foci in the presence of sera and foci at the highest dilution of respective sera sample). Each specimen was tested in two independent assays performed at different times. The FRNT-mNG_50_ titers were interpolated using a 4-parameter nonlinear regression in GraphPad Prism 8.4.3. Samples with an FRNT-mNG_50_ value that was below the limit of detection were plotted at 20.

#### Single-cell RNA-seq

##### Sample preparation, hashing, and pooling

Single-cell RNA-seq libraries were generated using the 10x Genomics Chromium 3’ Single Cell Gene Expression assay (#1000121) and Chromium Controller Instrument according to the manufacturer’s published protocol with modifications for cell hashing (Stoeckius et al., 2018). To block off-target antibody binding, Blocking Solution (5 μL of Human Trustain FcX (BioLegend #422302), and13.7 μL of a 10% Bovine Serum Albumin (BSA)) was added to 500,000 cells suspended in 50 μL Dulbecco’s Phosphate Buffered Saline (DPBS; Corning Life Sciences #21-031-CM) and incubated for 10 minutes on ice. To stain samples, 0.5 μg (1 μL) of a TotalSeq™-A anti-human Hashtag Antibody was suspended in 31.3 μL DPBS/2% BSA, then added to each sample with. For each batch of samples, 100,000 cells from 8 hashedsamples with a distinct Hashtag Antibody were pooled into Pool 1; 8 additional samples were pooled using the same method into Pool 2. Roughly 20,000 cells from a Leukopak healthy control were also labeled with a distinct TotalSeq™-A Hashtag Antibody, and were spiked into each pool to serve as a batch control.

##### Droplet encapsulation and reverse transcription

From each pool, 64,000 cells were loaded into each well of a Chromium Single Cell Chip G (10x Genomics #**1000073**) (8 wells per chip), targeting a recovery of 20,000 singlets from each well. Gel Beads-in-emulsion (GEMs) were then generated using the 10x Chromium Controller. The resulting GEM generation products were then transferred to semi-skirted 96-well plates and reverse transcribed on a C1000 Touch Thermal Cycler programmed at 53°C for 45 minutes, 85°C for 5 minutes, and a hold at 4°C. Following reverse transcription, GEMs were broken and the pooled single-stranded cDNA and Hashtag Oligo fractions were recovered using Silane magnetic beads (Dynabeads MyOne SILANE #37002D).

##### Library generation and separation

Barcoded, full-length cDNA including the Hashtag Oligos (HTOs) from the TotalSeq™-A Hashtag Antibodies were then amplified with a C1000 Touch Thermal Cycler programmed at 98°C for 3 minutes, 11 cycles of (98°C for 15 seconds, 63°C for 20 seconds, 72°C for 1 minute), 72°C for 1 minute, and a hold at 4°C. Amplified cDNA was purified and separated from amplified HTOs using a 0.6x size selection via SPRIselect magnetic bead (Beckman Coulter #22667) and a 1:10 dilution of the resulting cDNA was run on a Fragment Analyzer (Agilent Technologies #**5067-4626**) to assess cDNA quality and yield. HTO libraries were purified further with SPRIselect magnetic bead (Beckman Coulter #22667) and amplified and indexed with a custom HTO i7 index on a C1000 Touch Thermal Cycler programmed at 95°C for 3 minutes, 10 cycles of (95°C for 20 seconds, 64°C for 30 seconds, 72°C for 20 seconds), 72°C for 1 minute, and a hold at 4°C. The resulting HTO libraries were purified with SPRIselect magnetic bead (Beckman Coulter #22667) post-amplification and a 1:10 dilution of the resulting HTO libraries were run on a Fragment Analyzer (Agilent Technologies #**5067-4626**) to assess HTO quality and yield. A quarter of the cDNA sample (10 ul) was used as input for library preparation. Amplified cDNA was fragmented, end-repaired, and A-tailed is a single incubation protocol on a C1000 Touch Thermal Cycler programmed at 4°C start, 32°C for minutes, 65°C for 30 minutes, and a 4°C hold. Fragmented and A-tailed cDNA was purified by performing a dual-sided size-selection using SPRIselect magnetic beads (Beckman Coulter #22667). A partial TruSeq Read 2 primer sequence was ligated to the fragmented and A-tailed end of cDNA molecules via an incubation of 20°C for 15 minutes on a C1000 Touch Thermal Cycler. The ligation reaction was then cleaned using SPRIselect magnetic beads (Beckman Coulter #22667). PCR was then performed to amplify the library and add the P5 and indexed P7 ends (10x Genomics #1000084) on a C1000 Touch Thermal Cycler programmed at 98°C for 45 seconds, 13 cycles of (98°C for 20 seconds, 54°C for 30 seconds, 72°C for 20 seconds), 72°C for 1 minute, and a hold at 4°C. PCR products were purified by performing a dual-sided size-selection using SPRIselect magnetic beads (Beckman Coulter #22667) to produce final, sequencing-ready libraries.

##### Quantification and sequencing

Final libraries were quantified using Picogreen and their quality was assessed via capillary electrophoresis using the Agilent Fragment Analyzer HS DNA fragment kit and/or Agilent Bioanalyzer High Sensitivity chips. Libraries were sequenced on the Illumina NovaSeq platform using S4 flow cells. Read lengths were 28bp read1, 8bp i7 index read, 91bp read2.

#### Single-cell ATAC-seq

##### FACS neutrophil depletion

To remove dead cells, debris, and neutrophils prior to scATAC-seq as described previously (Swanson et al., 2021), PBMC samples were sorted by fluorescence activated cell sorting (FACS) prior to cell permeabilization. Cells were incubated with Fixable Viability Stain 510 (BD, 564406) for 15 minutes at room temperature and washed with AIM V medium (Gibco, 12055091) plus 25 mM HEPES before incubating with TruStain FcX (BioLegend, 422302) for 5 minutes on ice, followed by staining with mouse anti-human CD45 FITC (BioLegend, 304038) and mouse anti-human CD15 PE (BD, 562371) antibodies for 20 minutes on ice. Cells were washed with AIM V medium plus 25 mM HEPES and sorted on a BD FACSAria Fusion. A standard viable CD45+ cell gating scheme was employed; FSC-A x SSC-A (to exclude sub-cellular debris), two FSC-A doublet exclusion gates (FSC-W followed by FSC-H), dead cell exclusion gate (BV510 LIVE/DEAD negative), followed by CD45+ inclusion gate. Neutrophils (defined as SSC^high^, CD15^+^) were then excluded in the final sort gate. An aliquot of each post-sort population was used to collect 50,000 events to assess post-sort purity.

##### Sample preparation

Permeabilized-cell scATAC-seq was performed as described previously (Swanson et al., 2021). A 5% w/v digitonin stock was prepared by diluting powdered digitonin (MP Biomedicals, 0215948082) in DMSO (Fisher Scientific, D12345), which was stored in 20 μL aliquots at −20°C until use. To permeabilize, 1×10^6^ cells were added to a 1.5 mL low binding tube (Eppendorf, 022431021) and centrifuged (400×g for 5 min at 4°C) using a swinging bucket rotor (Beckman Coulter Avanti J-15RIVD with JS4.750 swinging bucket, B99516). Cells were resuspended in 100 μL cold isotonic Permeabilization Buffer (20 mM Tris-HCl pH 7.4, 150 mM NaCl, 3 mM MgCl2, 0.01% digitonin) by pipette-mixing 10 times, then incubated on ice for 5 min, after which they were diluted with 1 mL of isotonic Wash Buffer (20 mM Tris-HCl pH 7.4, 150 mM NaCl, 3 mM MgCl2) by pipette-mixing five times. Cells were centrifuged (400×g for 5 min at 4°C) using a swinging bucket rotor, and the supernatant was slowly removed using a vacuum aspirator pipette. Cells were resuspended in chilled TD1 buffer (Illumina, 15027866) by pipette-mixing to a target concentration of 2,300-10,000 cells per μL. Cells were filtered through 35 μm Falcon Cell Strainers (Corning, 352235) before counting on a Cellometer Spectrum Cell Counter (Nexcelom) using ViaStain acridine orange/propidium iodide solution (Nexcelom, C52-0106-5).

##### Tagmentation and fragment capture

scATAC-seq libraries were prepared according to the Chromium Single Cell ATAC v1.1 Reagent Kits User Guide (CG000209 Rev B) with several modifications. 15,000 cells were loaded into each tagmentation reaction. Permeabilized cells were brought up to a volume of 9 μl in TD1 buffer (Illumina, 15027866) and mixed with 6 μl of Illumina TDE1 Tn5 transposase (Illumina, 15027916). Transposition was performed by incubating the prepared reactions on a C1000 Touch thermal cycler with 96– Deep Well Reaction Module (Bio-Rad, 1851197) at 37°C for 60 minutes, followed by a brief hold at 4°C. A Chromium NextGEM Chip H (10x Genomics, 2000180) was placed in a Chromium Next GEM Secondary Holder (10x Genomics, 3000332) and 50% Glycerol (Teknova, G1798) was dispensed into all unused wells. A master mix composed of Barcoding Reagent B (10x Genomics, 2000194), Reducing Agent B (10x Genomics, 2000087), and Barcoding Enzyme (10x Genomics, 2000125) was then added to each sample well, pipette-mixed, and loaded into row 1 of the chip. Chromium Single Cell ATAC Gel Beads v1.1 (10x Genomics, 2000210) were vortexed for 30 seconds and loaded into row 2 of the chip, along with Partitioning Oil (10x Genomics, 2000190) in row 3. A 10x Gasket (10x Genomics, 370017) was placed over the chip and attached to the Secondary Holder. The chip was loaded into a Chromium Single Cell Controller instrument (10x Genomics, 120270) for GEM generation. At the completion of the run, GEMs were collected and linear amplification was performed on a C1000 Touch thermal cycler with 96–Deep Well Reaction Module: 72°C for 5 min, 98°C for 30 sec, 12 cycles of: 98°C for 10 sec, 59°C for 30 sec and 72°C for 1 min.

##### Sequencing library preparation

GEMs were separated into a biphasic mixture through addition of Recovery Agent (10x Genomics, 220016), the aqueous phase was retained and removed of barcoding reagents using Dynabead MyOne SILANE (10x Genomics, 2000048) and SPRIselect reagent (Beckman Coulter, B23318) bead clean-ups. Sequencing libraries were constructed by amplifying the barcoded ATAC fragments in a sample indexing PCR consisting of SI-PCR Primer B (10x Genomics, 2000128), Amp Mix (10x Genomics, 2000047) and Chromium i7 Sample Index Plate N, Set A (10x Genomics, 3000262) as described in the 10x scATAC User Guide. Amplification was performed in a C1000 Touch thermal cycler with 96–Deep Well Reaction Module: 98°C for 45 sec, for 9 to 11 cycles of: 98°C for 20 sec, 67°C for 30 sec, 72°C for 20 sec, with a final extension of 72°C for 1 min. Final libraries were prepared using a dual-sided SPRIselect size-selection cleanup. SPRIselect beads were mixed with completed PCR reactions at a ratio of 0.4x bead:sample and incubated at room temperature to bind large DNA fragments. Reactions were incubated on a magnet, the supernatant was transferred and mixed with additional SPRIselect reagent to a final ratio of 1.2x bead:sample (ratio includes first SPRI addition) and incubated at room temperature to bind ATAC fragments. Reactions were incubated on a magnet, the supernatant containing unbound PCR primers and reagents was discarded, and DNA bound SPRI beads were washed twice with 80% v/v ethanol. SPRI beads were resuspended in Buffer EB (Qiagen, 1014609), incubated on a magnet, and the supernatant was transferred resulting in final, sequencing-ready libraries.

##### Quantification and sequencing

Final libraries were quantified using a Quant-iT PicoGreen dsDNA Assay Kit (Thermo Fisher Scientific, P7589) on a SpectraMax iD3 (Molecular Devices). Library quality and average fragment size was assessed using a Bioanalyzer (Agilent, G2939A) High Sensitivity DNA chip (Agilent, 5067-4626). Libraries were sequenced on the Illumina NovaSeq platform with the following read lengths: 51nt read 1, 8nt i7 index, 16nt i5 index, 51nt read 2.

##### Extended PASC cohort for Olink analysis

To confirm some of the inflammatory signals observed in the cohort of 19 individuals (including 5 PASC), we selected additional PASC individuals from the longitudinal cohort regardless of their availability of serum collections within 15 days of diagnosis. We also included additional follow-up visits for the original PASC participants. All pre-vaccine visits from 55 PASC who had confirmed symptoms for more than 60 days PSO were included in the extended analysis. As controls, a group of 12 non-PASC recovered individuals were included in the analysis and they were selected for having: 1) at least a blood draw collected, 2) follow-up for at least 60 days to confirm their recovery, and 3) median age and age ranges comparable to the PASC individuals.

When combined with the cohort of 19 individuals described in Figure 1, PASC and Recovered individuals had the following characteristics:

**Table:** Olink Cohort Demographics and Baseline Characteristics for Symptomatic, PCR+ Individuals.

| Characteristic | PASC<br>(N=60) | Recovered<br>(N=26) | Total<br>(N=86) |
| --- | --- | --- | --- |
| Age, median (range)— years | 49.1 (21.7-82.1) | 48.6 (20.2-79.7) | 49.1 (20.2-82.1) |
| **Female sex at birth— no. (%)** | 40 (66.7) | 15 (57.7) | 55 (64.0) |
| **Race or ethnic group— no. (%)** |  |  |  |
| **White** | 56 (93.3) | 24 (92.3) | 80 (93.0) |
| **Hispanic or Latino** | 5 (8.3) | 2 (7.7) | 7 (8.1) |
| **Black or African American** | 0 (0.0) | 0 (0.0) | 0 (0.0) |
| **Asian** | 0 (0.0) | 1 (3.8) | 1 (1.2) |
| **Other<sup>a</sup>** | 4 (6.7) | 1 (3.8) | 5 (5.8) |
| **Median acute symptom duration (range)— days** | 17.5 (2-94) | 11.0 (1-24) | 15 (1-94) |
| **Disease severity (WHO Score)<sup>b</sup> —no. (%)** |  |  |  |
| **2 (Mild)** | 32 (53.3) | 18 (69.2) | 50 (58.8) |
| **3 (Moderate)** | 25 (41.7) | 7 (26.9) | 32 (37.6) |
| **5 (Severe)** | 3 (5.0) | 0 (0.0) | 3 (3.5) |
| **Median cumulative illness severity score (range)** | 29.5 (3-233) | 13.0 (0-49) | 23.5 (0-233) |
| **Median of maximum days of per-participant follow-up (range)** | 236.5 (21-379) | 104 (32-354) | 201 (21-379) |

<sup>a</sup>Individuals identifying as Other included: Other (1); White; Other (n=1); American Indian or Alaska Native; White (n=1); Asian, Black or African American (n=1); Asian; White (n=1); <sup>b</sup>1 non-PASC recovered participant asymptomatic with no WHO score.

#### Olink serum protein measurement

Serum samples were inactivated with 1% Triton X-100 for 2h at room temperature according to the Olink COVID-19 inactivation protocol. Inactivated samples were then run on the Olink Explore 1536 platform, which uses paired antibody proximity extension assays (PEA) and a next generation sequencing (NGS) readout to measure the relative expression of 1472 protein analytes per sample. Analytes from the inflammation, oncology, cardiometabolic, and neurology panels were measured.

For plate setup, samples were randomized across plates to achieve a balanced distribution of age and gender. Longitudinal samples from the same participant were run on the same plate. To facilitate comparisons with future batches, sera from 15 donors was commercially purchased (BioIVT) and randomly interspersed amongst the above study samples. Commercial samples included serum from COVID-19 serology-negative, serology-positive, PCR-positive, and recovered (no longer symptomatic) participants.

Data were first normalized to an extension control that was included in each sample well. Plates were then standardized by normalizing to inter-plate controls run in triplicate on each plate. Data were then intensity normalized across all samples. Final normalized relative protein quantities were reported as log2 normalized protein expression (NPX) values.

#### Olink preprocessing

Olink results and QC flags were reviewed for overall quality. Results for TNF, IL6 and CXCL8, which were measured on all 4 Olink panels, were reviewed prior to averaging to a single NPX value for analysis. Two samples had discrepant cross-panel measurements on these proteins. The results that trended most consistently with the participant’s longitudinal measurements were kept and averaged. Serum samples were analyzed in two batches. Following the method recommended by Olink, results of the later batch were bridged to those of the earlier batch using a set of 42 cohort samples that were tested in both batches. A batch offset for each analyte was calculated as the median difference on the 42 samples as measured between the two batches, excluding samples with QC warning flags. The analyte-specific offsets were then added to the raw NPX values of the later batch.

### Data analyses

#### Illness severity metrics and scoring

For acute COVID-19, categorization of illness severity was classified by participant report of impact on Activities of Daily Living (ADLs) for each day of illness (U S Department et al., 2017). Severe days hospitalized were also recorded as were any treatment or therapies received. Participants were scored according to their maximum severity for each day: 0, no symptoms; 1, mild impact on ADLs reported; 2, moderate impact on ADLs reported; 3, severe illness without hospitalization; 4, severe illness with hospitalization; 5, life threatening illness hospitalized with ICU care. Durations were assigned for days spent at each illness severity. A cumulative illness severity score was calculated for each participant by multiplying each severity score by the number of days spent at each level, then summing all values. Participants were classified as post-acute sequelae of SARS-CoV-2 infection (PASC) if any symptoms continued from acute illness or related to COVID-19 beyond 60 days. Sickest PASC participants had ongoing symptoms that significantly impacted activities of daily living, quality of life, and inability to continue employment.

#### Metadata correlations

Spearman’s rank correlation coefficients were calculated for pairwise combinations of SARS-CoV-2-specific immune response estimates and clinical metadata. Symptoms at diagnosis and comorbidities with ≤ 3 events were excluded from analysis. P-values were adjusted using Benjamini-Hochberg method and adjusted p-values ≤ 0.05 were considered significant.

#### Flow cytometry

Following sample acquisition on the Cytek Aurora flow cytometer, samples were unmixed using library reference controls pre-recorded on the instrument using the batch control sample, and the data files were exported in FCS 3.1 format. Unmixing performance was visually checked and unmixing errors were adjusted with compensation on the batch control sample in each individual batch using FlowJo v10.7.1. Compensation adjustments from the batch control were applied to the entire batch. The fluorescent channels in each compensated FCS file were transformed using the logicle transform. The compensated and transformed data was analyzed using bivariate hierarchical gating. Initial placement of gates was implemented automatically using the openCyto R package using customized gating templates for each panel. Following the preliminary automated gate placement, all gates were visually checked for errors and corrected manually per sample in a Flowjo v10.7.1 workspace file generated with the CytoML R package (Finak et al., 2018). Gating plots for all samples from each participant were generated with the ggCyto R package (Van et al., 2018), and gates were further reviewed for consistency and batch effects across longitudinal timepoints. For each population gate, a cutoff of 50 events was also set, below which populations were not reported. Proportions for each cell subtype were calculated by dividing the counts for the gated subpopulation by the total number of CD45+ leukocytes for each sample. Proportions of activated cell populations were calculated by dividing the counts for each positive activation marker population by the counts of the corresponding parent population. Samples were run in multiple batches with bridging controls and we determined that the proportion of variability due to batch was lower than biological variability across samples (**Fig. S1b**).

#### Participant-specific linear modeling of binding antibodies, intercept and slope

Linear mixed effects models were used to estimate binding antibody RBD titer, *Y_ij_*, as a function of *t_ij_*, the *j^th^* time since symptom onset for the *i^th^* individual, with random effects for intercept and slope and *t_ij_* > 30 days for all *i*, *j*:

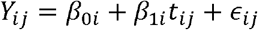

where *β*_0*i*_ = *β_0_* + *b_i_* and *β*_1*i*_ = *β_1_* + *c_i_* with (*b_i_*, *c_i_*) iid ~ *N*_2_ (0, *Σ*), with

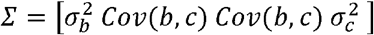

and 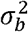 and 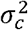 are the between-person variation in the intercept and slope of log RBD titer responses respectively, *Cov(b, c)* is the covariance between the intercept and slope, and *ϵ_ij_* iid ~ *N*(0, σ^2^). The random effects, *b_i_* and *c_i_*, are each assumed to be independent for different individuals and the within-individual errors *ϵ_ij_* are assumed to be independent for different *i, j* and to be independent of the random effects. A similar model was fit to the untransformed IgG N indices, which are linear on the original scale. The function lme from the R package nlme was used to fit the models. Individual-level estimates of the intercepts, slopes, day 30 and day 180 were obtained from the models.

#### scRNA-seq analysis

##### Data preprocessing

Binary Base Call (BCL) files were demultiplexed using cellranger mkfastq (10x Genomics v3.0.2) to produce FASTQ files for scRNA-seq and HTO barcode libraries. scRNA-seq FASTQ files for each hashed well were aligned to the 10x Genomics GRCh38 reference transcriptome (10x Genomics vGRCh38-3.0.0) using cellranger (10x Genomics v3.1.1) using default settings. HTO barcode FASTQ files were processed using BarCounter (Swanson et al., 2021) to quantify the count of each HTO barcode for each cell barcode. A custom R pipeline, BarcodeTender (to be released at https://github.com/AllenInstitute/BarcodeTender-pipeline), was used to deconstruct gene-by-cell matrices from cellranger outputs and assign cells to their originating participant sample based on HTO counts. For each well, a threshold was determined for each HTO by removing barcodes with low counts, then performing 1 dimensional k-means clustering to group cells into a “high” or “low” group. The minimum count for the “high” group was set as a cutoff, and all cells above that cutoff were considered positive for that HTO barcode. If the mean of the “high” group was not greater than 4-fold of the mean of the “low” group, the minimum cutoff was updated to the mean of the “low” group, and the process was iterated. If a separation could not be established, the cutoff was set to the maximum count, and all cells were considered negative for the HTO barcode. Cutoffs were then used to binarize cell barcodes as positive or negative for each HTO, and the number of positive HTOs was calculated for each cell barcode. Cell barcodes positive for only one HTO barcode were considered “singlets”, and any cells with two or more were considered “multiplets”. For each well, count matrices generated by cellranger were split to select singlet cell barcodes from each PBMC sample based on HTO barcodes, then cells from each PBMC sample were concatenated into separate matrices for each participant. Throughout the process, metadata tracking the originating Batch and 10x Chromium Chip and Well were retained to enable batch effect analysis. During the splitting and merging process, quality control reports were generated to allow review prior to downstream analysis.

##### Cell type labeling

We utilized the supervised PCA projection and anchor-based transfer approach implemented in Seurat v4.0.0 to map cells to a reference projection generated using the Weighted Nearest Neighbor (WNN) graph algorithm, as described in (Hao et al., 2020b). A CITE-seq dataset from 8 healthy HIV vaccine volunteers during the course of vaccination was used as the reference dataset to label cells. The CITE-seq reference consisted of 228 surface proteins (antibody-derived tags, ADT) and approximately 160k single cells. The reference consisted of three levels of cell label granularity (Levels 1, 2 and 3). We constructed a level 2.5 label structure that consists of all cell type annotations from level 2 except merging the “CD8 TEM_4” and “CD8 TEM_5” from level 2, into a “CD8 TEMRA” label (based on marker gene expression) and replacing the “Tregs” label in level 2, with “Treg naive” and “Treg memory” labels from level 3, thus spanning a complete spectrum of cell type annotations. We assessed the distribution of label transfer scores per cell and retained cells with a label score ≥0.5. Batch variability was lower than biological variation contributed by cell type and participant-specific differences (**Fig. S1d, S1e**).

##### Differential expression early acute infection visit compared to uninfected controls

The hurdle model implemented in the MAST package (Finak et al., 2015) was used to identify differentially expressed genes between the early acute infection COVID-19 participants compared to uninfected controls. An age matched comparison between infected and uninfected controls was performed, i.e older (> 40 years) COVID-19 participants versus older uninfected controls and younger (< 40 years) COVID-19 participants versus younger uninfected controls. Genes that were expressed in at least 10% of all cells were considered for downstream analysis. Gene expression was normalized by scaling raw counts by the total number of reads per cell multiplied by a scaling factor of 10,000, then using a log2 transformation. A hurdle model was fit on the filtered and normalized data, modeling the infection status and adjusting for the batch for every cell type. A likelihood ratio test was then performed to assess if the coefficients are different from zero. The p-values were adjusted for multiple comparisons using the Benjamini and Hochberg (BH) method (Benjamini and Hochberg, 1995). Adjusted p-values < 0.05 were considered significant.

##### Participant-specific longitudinal gene expression changes

Longitudinal changes in gene expression were also identified using the hurdle model implemented in the MAST package. A hurdle model was fit to each COVID-19 participant independently in order to identify participant-specific longitudinal transcriptomic changes. Genes that were expressed in at least 10% of cells per participant were considered for this analysis. The models were fit on the filtered and normalized data, modeling the days since symptom onset as a continuous variable within each cell type and adjusting for the batch only if any timepoints from the same participant were run across multiple batches. A likelihood ratio test was then performed to assess if the coefficients are different from zero. Obtained p-values are adjusted for multiple comparisons using the BH method. Adjusted p-values < 0.05 were considered significant.

##### Pathway enrichment analysis

Gene Set Enrichment Analysis (GSEA) (Subramanian et al., 2005) was performed among genes that defined early acute infection status and genes that defined longitudinal changes. A custom collection of genesets that included the Hallmark v7.2 genesets, KEGG v7.2 and Reactome v7.2 from the Molecular Signatures Database (MSigDB, v4.0) was used as the pathway database. The “Type III interferon signaling” gene set was manually curated from the Interferome database (Rusinova et al., 2013). Genes were pre-ranked by the decreasing order of their log fold changes or coefficients. The running sum statistics and Normalized Enrichment Scores (NES) were calculated for each comparison. The pathway enrichment p-values were adjusted using the BH method and pathways with p-values < 0.05 were considered significantly enriched.

##### Sample-level enrichment (SLEA)

Sample-level enrichment analysis (SLEA, (Gundem and Lopez-Bigas, 2012) was used to represent the GSEA pathway expression results on a per-sample basis. The SLEA score was calculated by first calculating the mean expression value of genes (averaged across single cells) enriched in a pathway, then comparing it to the mean expression of random sets of genes (averaged across single cells) of the same size for 1,000 permutations per sample. The difference between the observed and expected mean expression values for each pathway was determined as the SLEA pathway score per sample.

#### scATAC-seq analysis

##### Data preprocessing

scATAC-seq libraries were processed as described previously (Swanson et al., 2021). In brief, cellranger-atac mkfastq (10x Genomics v1.1.0) was used to demultiplex BCL files to FASTQ. FASTQ files were aligned to the human genome (10x Genomics refdata-cellranger-atac-GRCh38-1.1.0) using cellranger-atac count (10x Genomics v1.1.0) with default settings. Fragment positions were used to quantify reads overlapping a reference peak set (GSE123577_pbmc_peaks.bed.gz from GEO accession GSE123577; (Lareau et al., 2019)) which was converted from hg19 to hg38 using the liftOver package for R (Lawrence et al., 2009), ENCODE reference accessible regions (ENCODE file ID ENCFF503GCK; (Vierstra et al., 2020)), and TSS regions (TSS ±2kb from Ensembl v93; (Yates et al., 2020)) for each cell barcode using a bedtools (v2.29.1; (Quinlan and Hall, 2010)) analytical pipeline.

##### Quality Control

Custom R scripts were used to remove cells with less than 1,000 uniquely aligned fragments, less than 20% of fragments overlapping reference peak regions, less than 20% of fragments overlappingENCODE TSS regions, and less than 50% of peaks overlapping ENCODE reference regions. The ArchR package (Granja et al., 2021) was used to assess doublets in scATAC data. Doublets were identified using the ScoreDoublets function using a filter ratio of 8, and cells with DoubletEnrichment scores from 0-1.16 were considered “singlets” and retained for further analysis. Samples with particularly high doublet scores across all cells (>70% of cells with DoubletEnrichment scores > 1.5) were not considered for downstream analysis.

##### Dimensionality reduction and cell type labeling

We used the ArchR package to generate a count matrix for the PBMC reference peak set described above (Lareau, et al. 2019 Nat Biotech). Dimensionality reduction was performed using the ArchR addIterativeLSI function (parameters varFeatures = 10000, iterations = 2), and the addClusters function was used to identify clusters in LSI dimensions using the Louvain community detection algorithm. For visualization, UMAP was performed using ArchR’s addUMAP function with default settings. The ArchR addGeneIntegrationMatrix function (parameters transferParams = list(dims = 1:10, k.weight = 20) was used to label our scATAC cells using the Seurat level 1 cell types from the Seurat v4.0 PBMC reference dataset (Hao et al., 2020b). We observed that Louvain clusters contained cells with mixed level 1 identity assignments from label transfer, and cluster labels were often spread across the UMAP space. In comparison, cell type assignments in UMAP coordinates seemed to cleanly separate cell-type identities. To generate clusters that more closely matched label transfer results, we performed K-means clustering on the UMAP coordinates using a range of number of cluster centers from 3 to 50, and identified a set of K-means clusters that each had > 80% of cells sharing a single cell type identity. Almost all such clusters contained >= 98% cells from a single major cell type class (T Cells, B Cells, NKs, or Monocytes/DCs/other), with the exception of a single cluster with 88% purity. We used K-means clusters that shared cell class identities to subset the data into T Cells, B Cells, NKs, or monocytes/DC/other classes for downstream analyses. For each broad type, we performed dimensionality reduction by Iterative LSI using 500 bp genomic tiles. We then performed a second round of label transfer using the ArchR addGeneIntegrationMatrix function (parameters as described for level 1, above) using the higher resolution level 2.5 cell identities (described for scRNA-seq label transfer, above) from the Seurat PBMC reference dataset.

##### Peak calling and motif enrichment analysis

For each of the broad types above, we grouped cells within each level 2.5 cell type label by participant, and used the ArchR addGroupCoverages and addReproduciblePeakSets functions to perform *de novo* peak calling to identify putative regulatory regions throughout the genome. We ensured that pseudo-bulk replicates derived from participant and cell type intersections were robust by requiring a minimum of 100 cells per pseudo-bulk replicate, with a minimum of 2 replicates for each group.. After identifying *de novo* peak sets, we annotated transcription factor binding sites (TFBS) in all peaks using the ArchR addArchRAnnotations function (parameter collection = “EncodeTFBS”) to label binding sites that have been previously observed in ENCODE datasets. We then used the ArchR addBgdPeaks and addDeviationsMatrix functions (parameter peakAnnotation = “EncodeTFBS”), based on ChromVar (Schep et al., 2017), to generate a measure of increased or decreased binding site accessibility compared to a random GC-matched peakset. Differential TFBS usage was computed using the ArchR getMarkerFeatures function (parameters bias = c(“TSSEnrichment”, “log10(nFrags)”), testMethod = “wilcoxon”) to identify differentially accessible genomic regions, followed by the peakAnnoEnrichment function (parameters peakAnnotation = “EncodeTFBS”, cutOff = “FDR <= 0.1”) to score differentially enriched TFBS deviations. Longitudinal TFBS usage scores were calculated by taking the mean TFBS deviation Z-score for each TF, participant, and time point among COVID19-positive samples.

#### Linear regression models of early acute infection

Linear regression models were fit to assess changes in normalized protein expression (NPX) for each protein from Olink, as well as flow cytometry or scRNA-seq cell proportions as a function of infection status using the *lm* function from the package *stats* in R. Participant age and biological sex were used as fixed effects in the model in order to control for these potentially confounding variables. P-values were adjusted using the BH method to control false discovery rate (FDR). Adjusted p-values < 0.05 were considered significant.

#### Linear mixed-effects models of longitudinal infection

Linear mixed-effects models (LME) were fit to assess the longitudinal changes in the serum proteome, flow cytometry and scRNA based cell type proportions as a function of days since symptom onset using the *lme* function (lme4 v1.1-26) implemented in R. Individual NPX protein expression and cell type proportion values were treated as dependent variables. However, COVID-19 participants appear to vary in the slopes and intercepts of days since symptoms. Thus, random effects for both the slope (days since symptom onset) and intercept (participant ID), and fixed effects for age and sex were used in the mixed effects model. P-values were calculated using Wald chi-square tests and adjusted using the BH method to control FDR. Adjusted p-values < 0.05 were considered significant.

Pathways enriched among significant longitudinally changing proteins were identified using Fisher’s over-representation analysis. Pathway p-values were adjusted for multiple comparisons using the BH method and pathways significant at adjusted p-value < 0.05 were reported

#### Outlier analysis

We performed outlier analysis to identify differential proteins in each COVID19 participant. To identify outliers, we calculated mean expression and standard deviation (SD) of each protein in uninfected participants. The normalized protein expression in COVID-19 participants was compared with the average expression from uninfected participants. Outliers were defined as proteins with expression greater than mean ± 2SD (uninfected participants).

#### Supervised principal coordinate analysis

Supervised principal coordinate analysis (PCoA) was performed on Olink data using the R function *pcoa* from the ape package (v5.5, Paradis and Schliep, 2019, Bioinformatics). Euclidean distances between proteins that were significantly different between COVID-19 participants at Visit 1 and uninfected controls and proteins that significantly changed over time among COVID-19 participants based on the linear regression models and linear mixed effect models were used to define the distance matrix between samples. The distance of a particular sample to the centroid of uninfected controls was then defined as 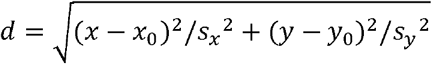, where *x, y* are the first and the second principal coordinates of the sample while *x*_o_, *y*_o_, *s*_x_, and *s*_y_ are the corresponding mean values and standard deviations of uninfected controls.

#### Integrative cell-cell network analysis

To identify the cell type specific ligand-receptor pairs significantly enriched in COVID-19 participants we selected scRNA-seq data from healthy participants and COVID-19 participant first visit samples for analysis using the nichnetr package (Browaeys et al., 2020). The ligand-target model retrieved from the nichnetr package includes 688 ligands and 25,345 potential downstream target genes, with values denoting the prior potential that a particular ligand might regulate the expression of a specific target gene. Receivers and senders were identified per cell type for plasmablasts, proliferating CD4+ T cells, proliferating CD8+ T cells, CD14+ monocytes, CD16+ monocytes, NK cells, proliferating NK cells, cDC1, cDC2, and pDC types. Genes were considered for downstream analysis if expressed in at least 10% of the cells from the corresponding cell type. Background gene expression data was obtained from samples of uninfected control participants. Ligand-receptor scores were ranked based on Pearson correlation coefficients (PCC). The top 50 identified ligands were then used to infer putatively active ligand-target links using the nichenetr get_weighted_ligand_target_links function with default parameters. The top 10 ligands and their targets were used to visualize the cell type-specific interaction network.

To calculate the changes in the ligand-receptor network as a result of COVID-19 infection, we identified differentially expressed genes from 2 comparisons: 1) acute infection (≤15 days PSO) compared to uninfected controls, and 2) all COVID-19 participant samples compared to uninfected controls (adjusted p-value < 0.05). All proteomic features were included for corresponding change in proteomic features.

## KEY RESOURCES TABLE

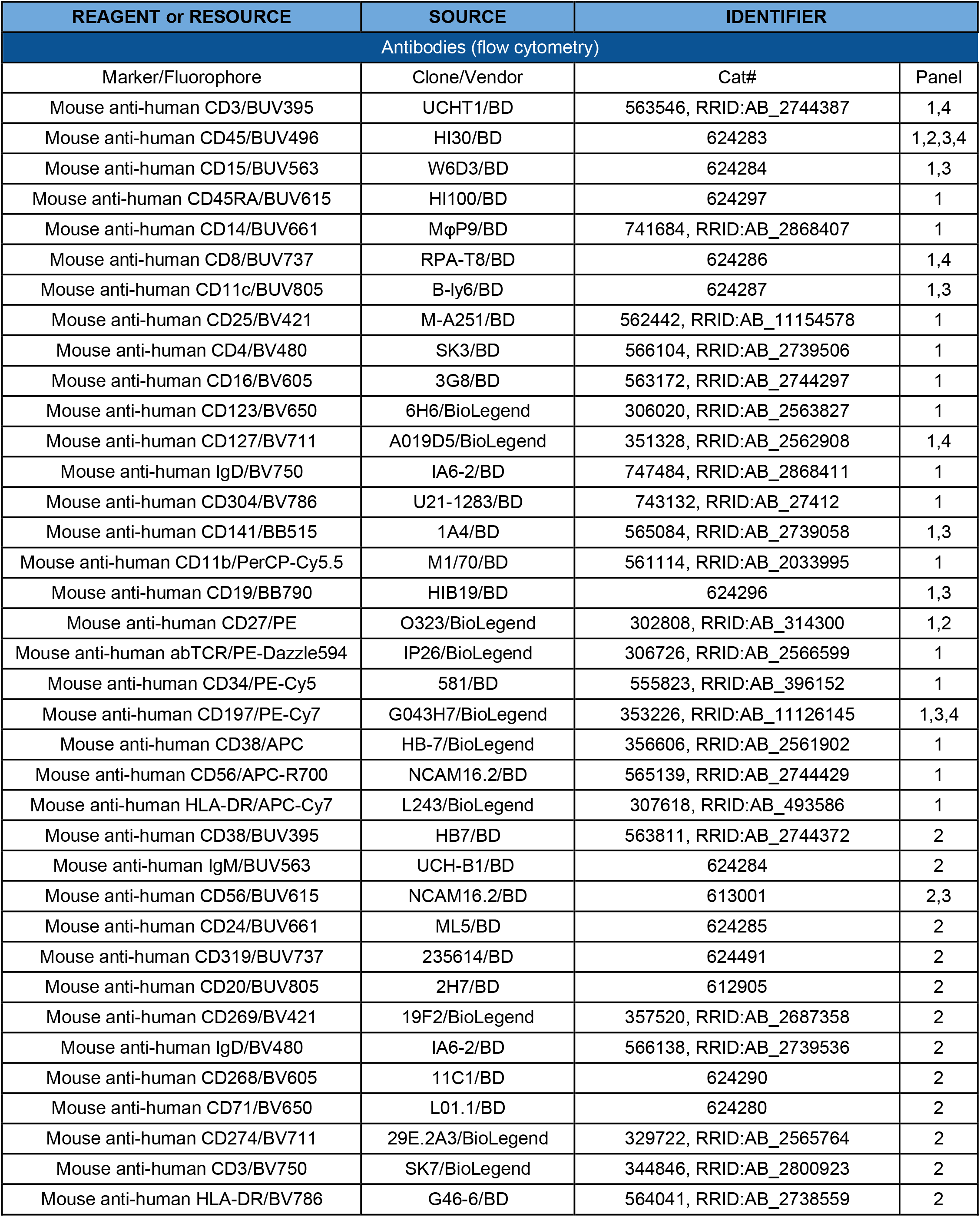
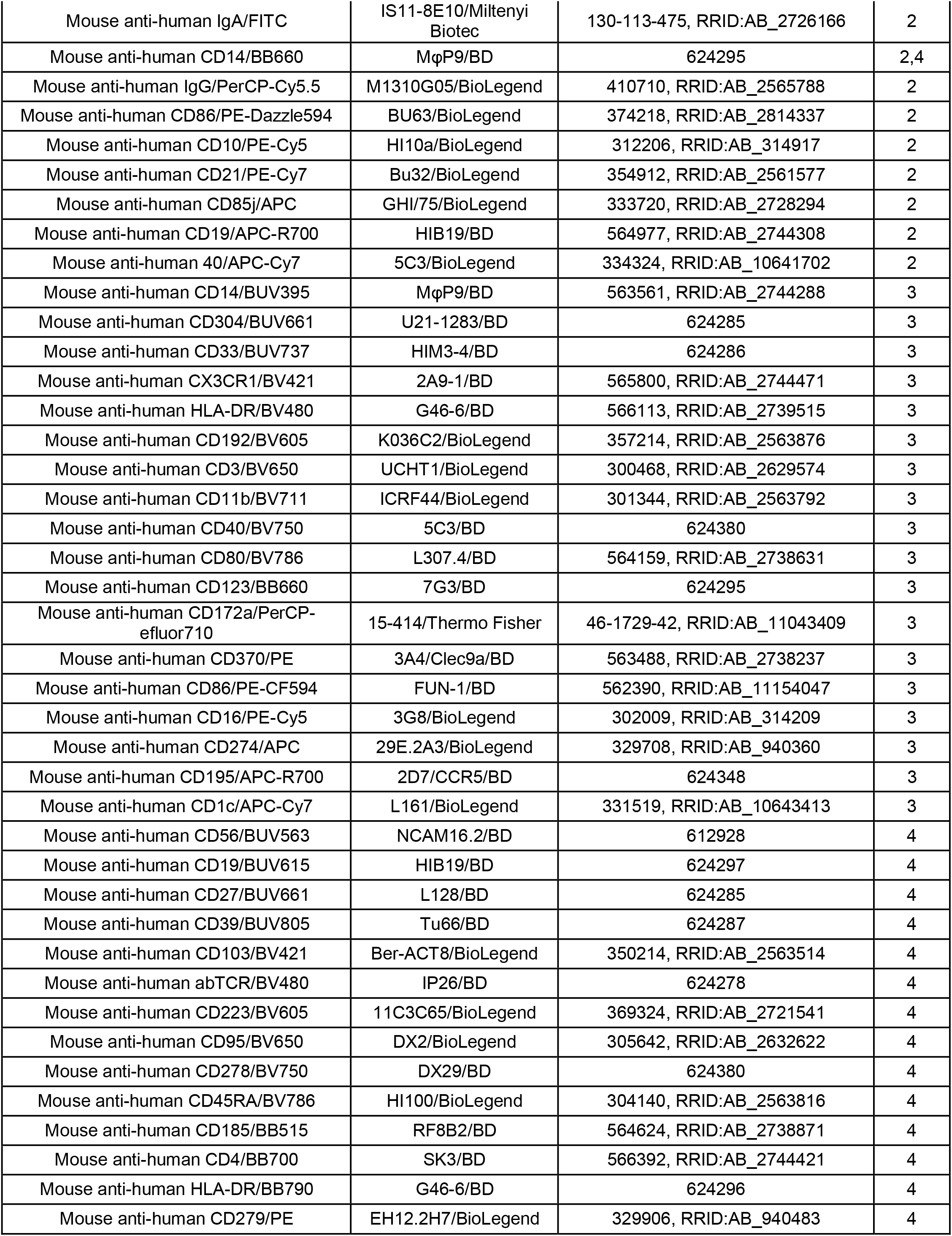
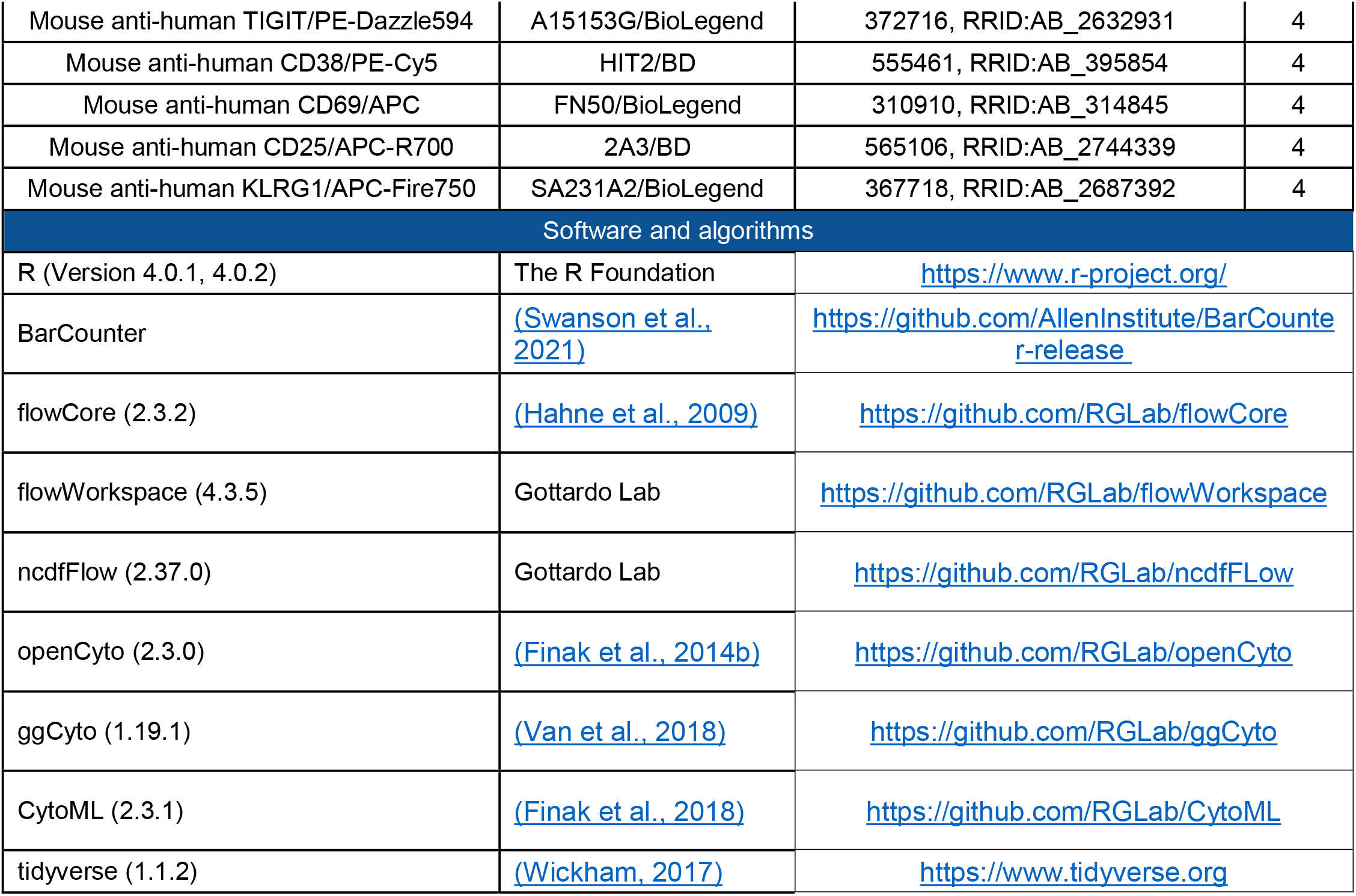

### Panels

1: PS1 (Survey)

2: PB1 (B cell)

3: PM1 (Myeloid/dendritic cell)

4: PT1 (T cell)

### Note

BD antibodies with a 600000 cat # are custom conjugates.

| <u>Lot Number(s):</u> | <u>Gender:</u> | <u>Age:</u> | <u>Race:</u> | <u># of Vials:</u> | <u>Target Concentration (per Vial):</u> | <u>Post-Processing Concentration (per Vial):</u> | <u>% Viability:</u> |
| --- | --- | --- | --- | --- | --- | --- | --- |
| HMN169517 | Male | 24 | Black | 330 | 10 X 10<sup>6</sup> | 16.2 x 10<sup>6</sup> | 94.9% |

