## Supplementary figures and images for "Longitudinal immune dynamics of mild COVID-19 define signatures of recovery and persistence"

### Supplemental Figure S1

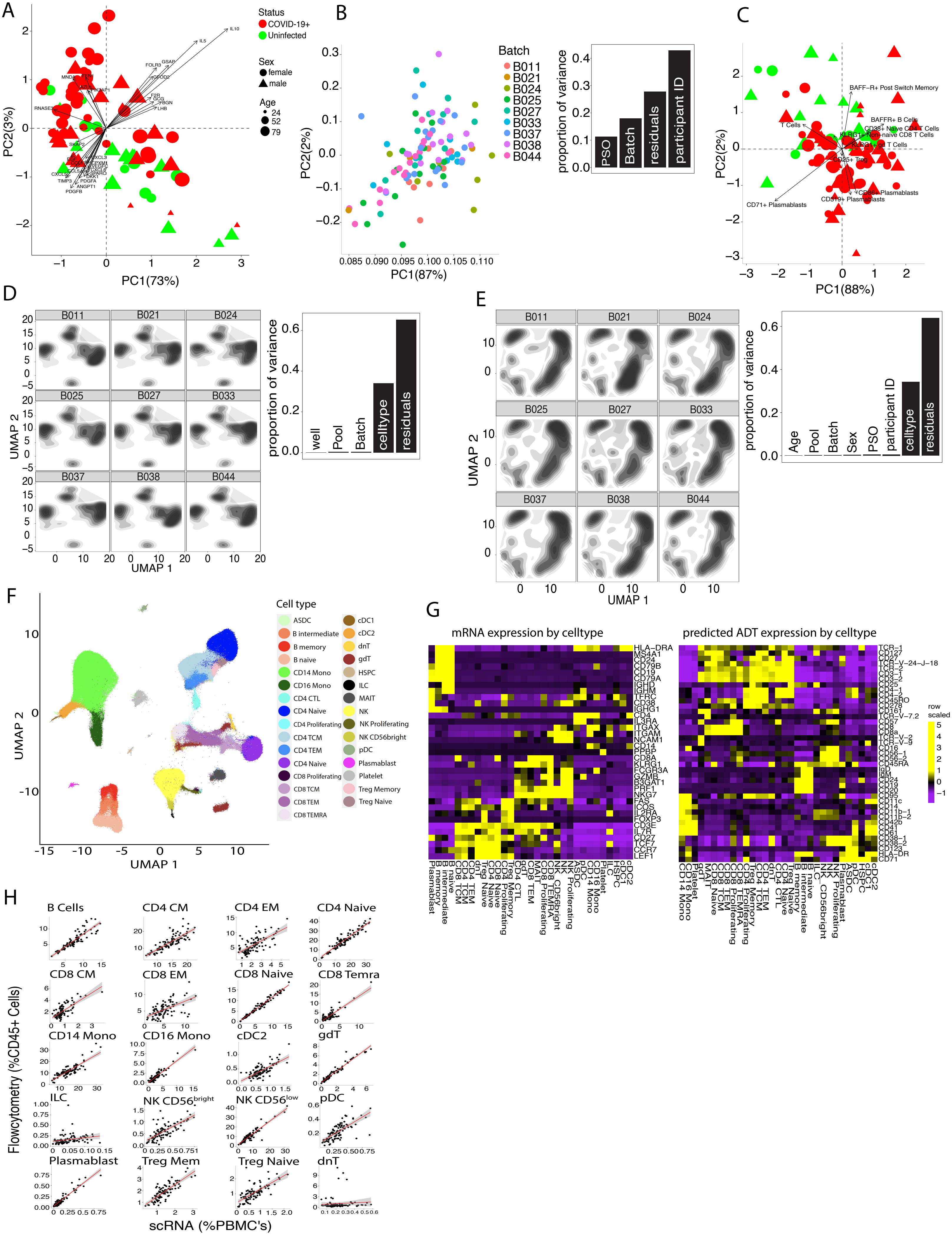

### Supplemental Figure S2

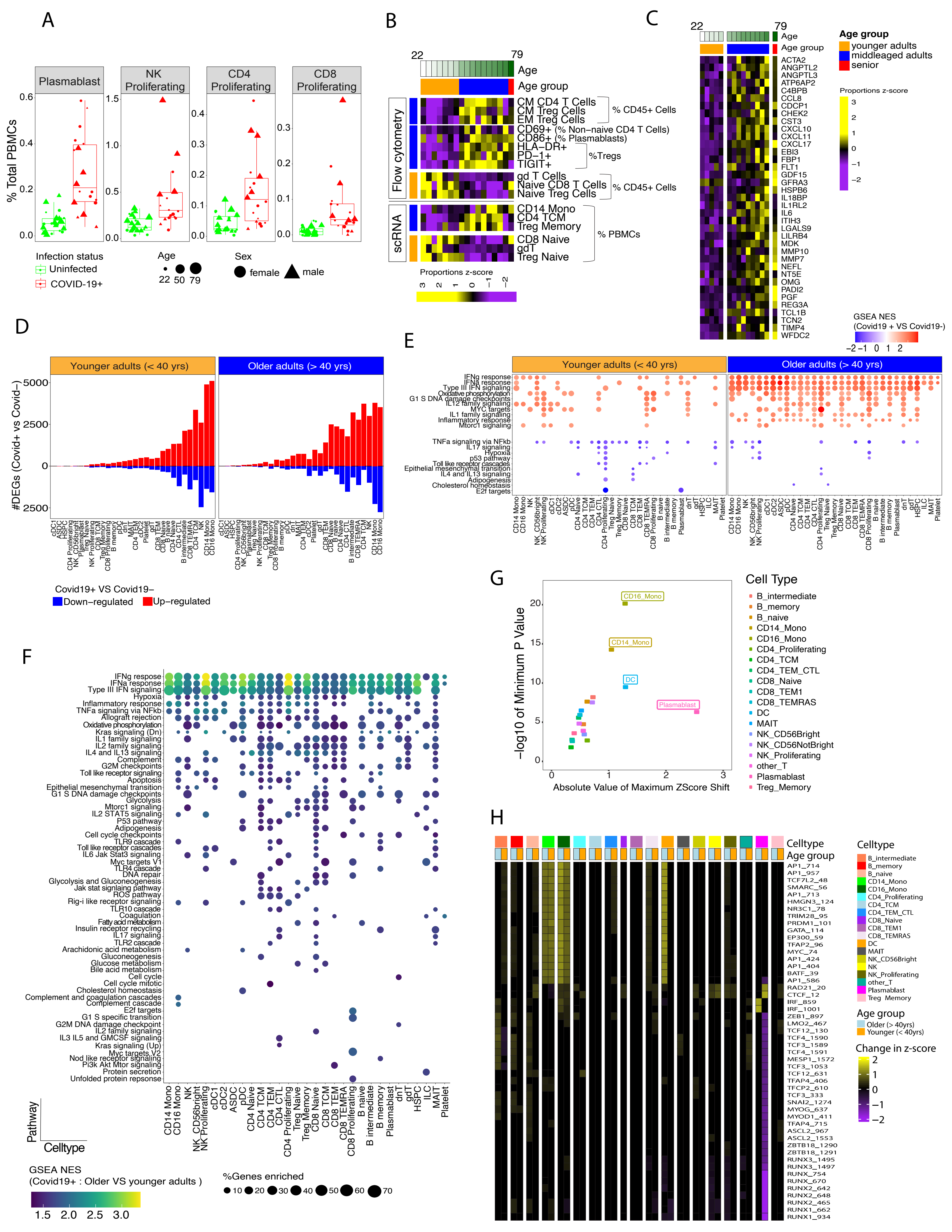

### Supplemental Figure S3

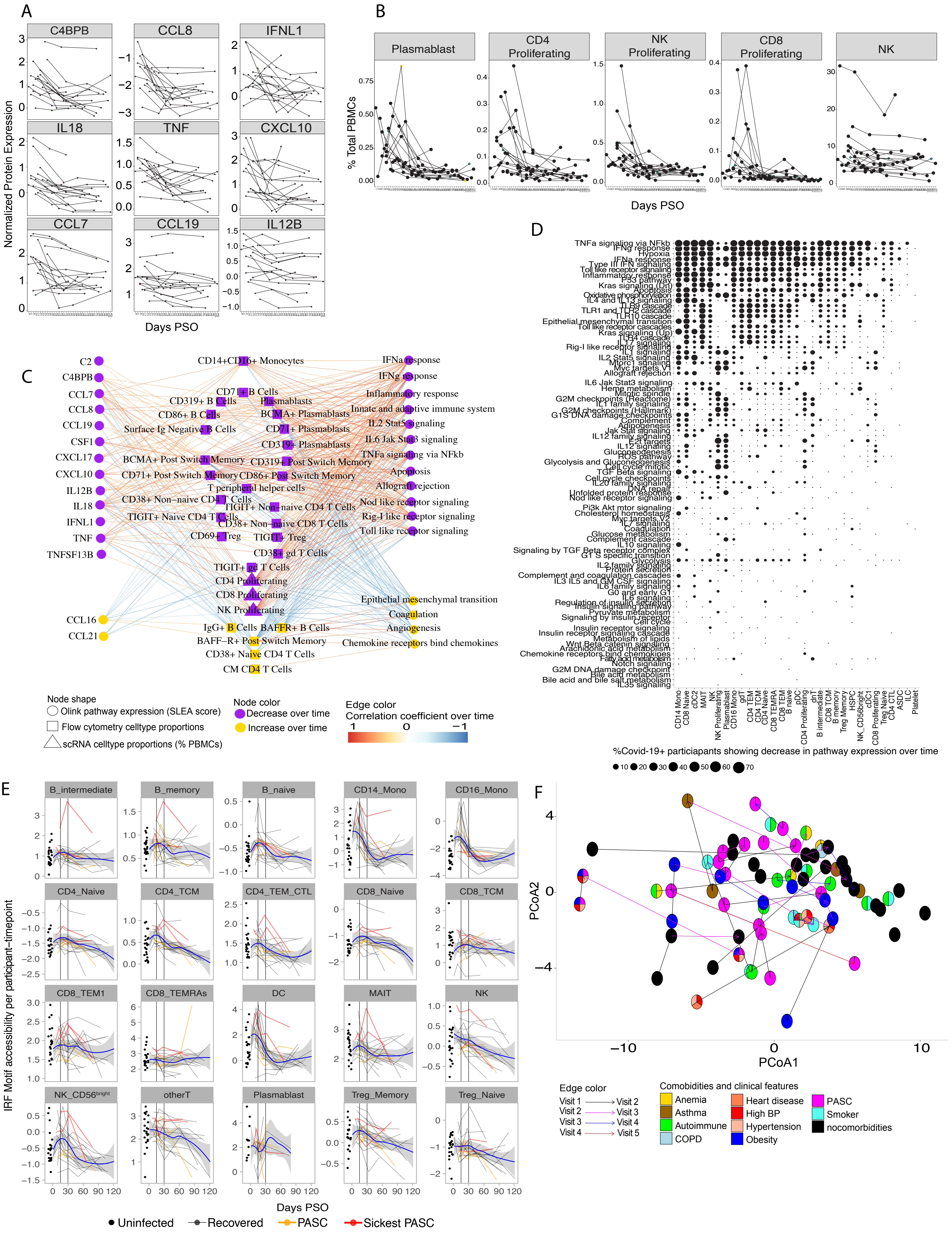

### Supplemental Figure S4

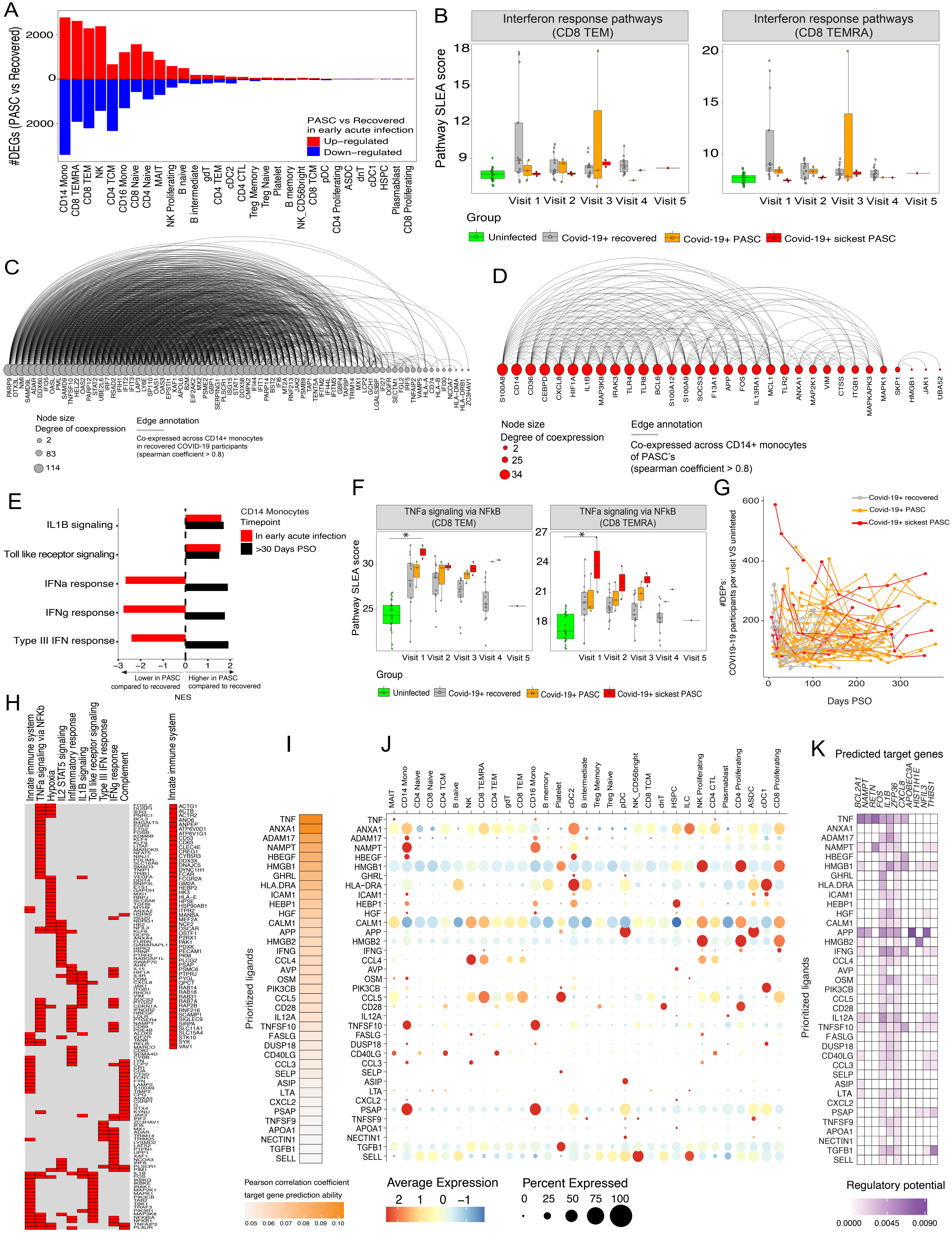

### Supplemental Figure S5

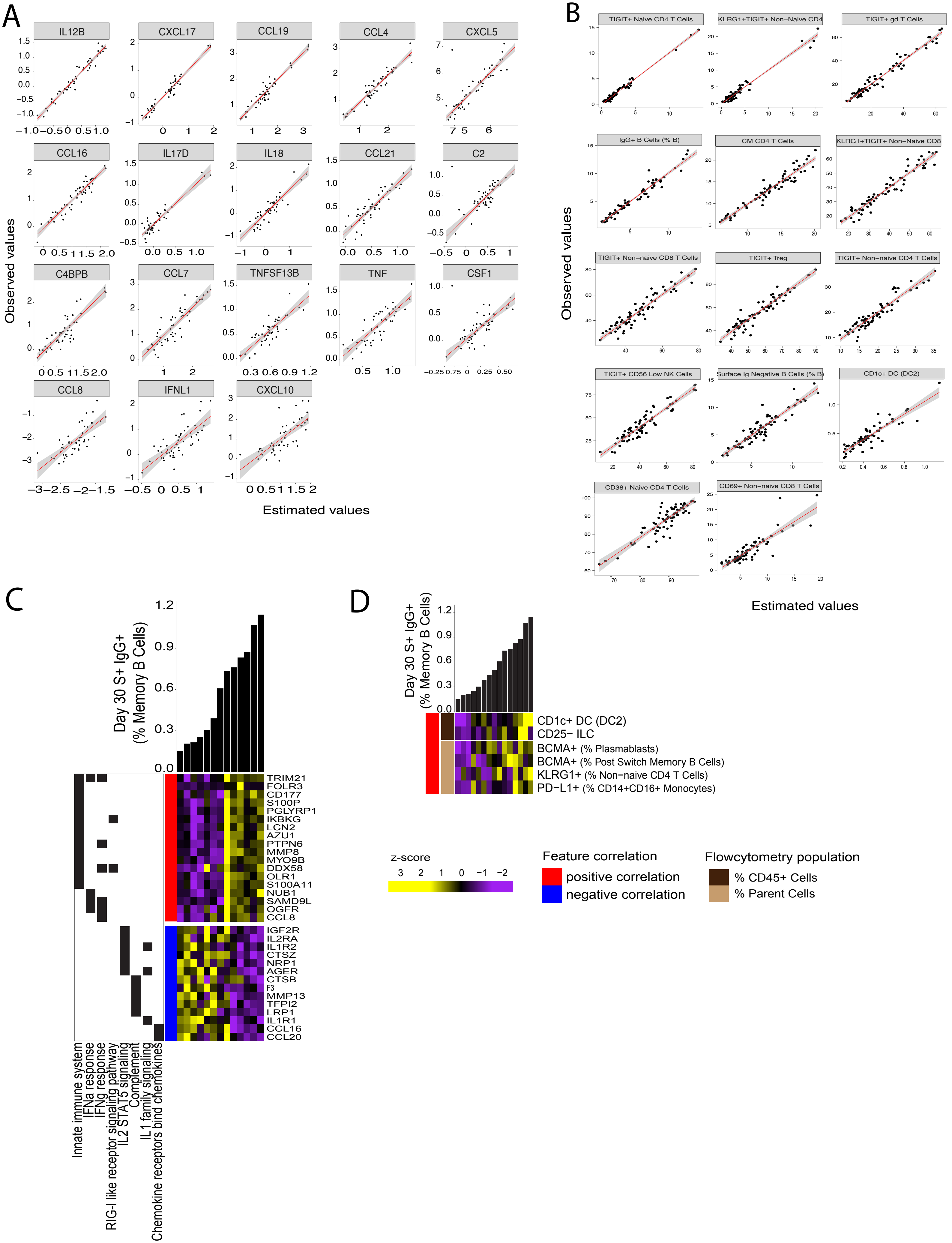

### Supplemental Figure S6

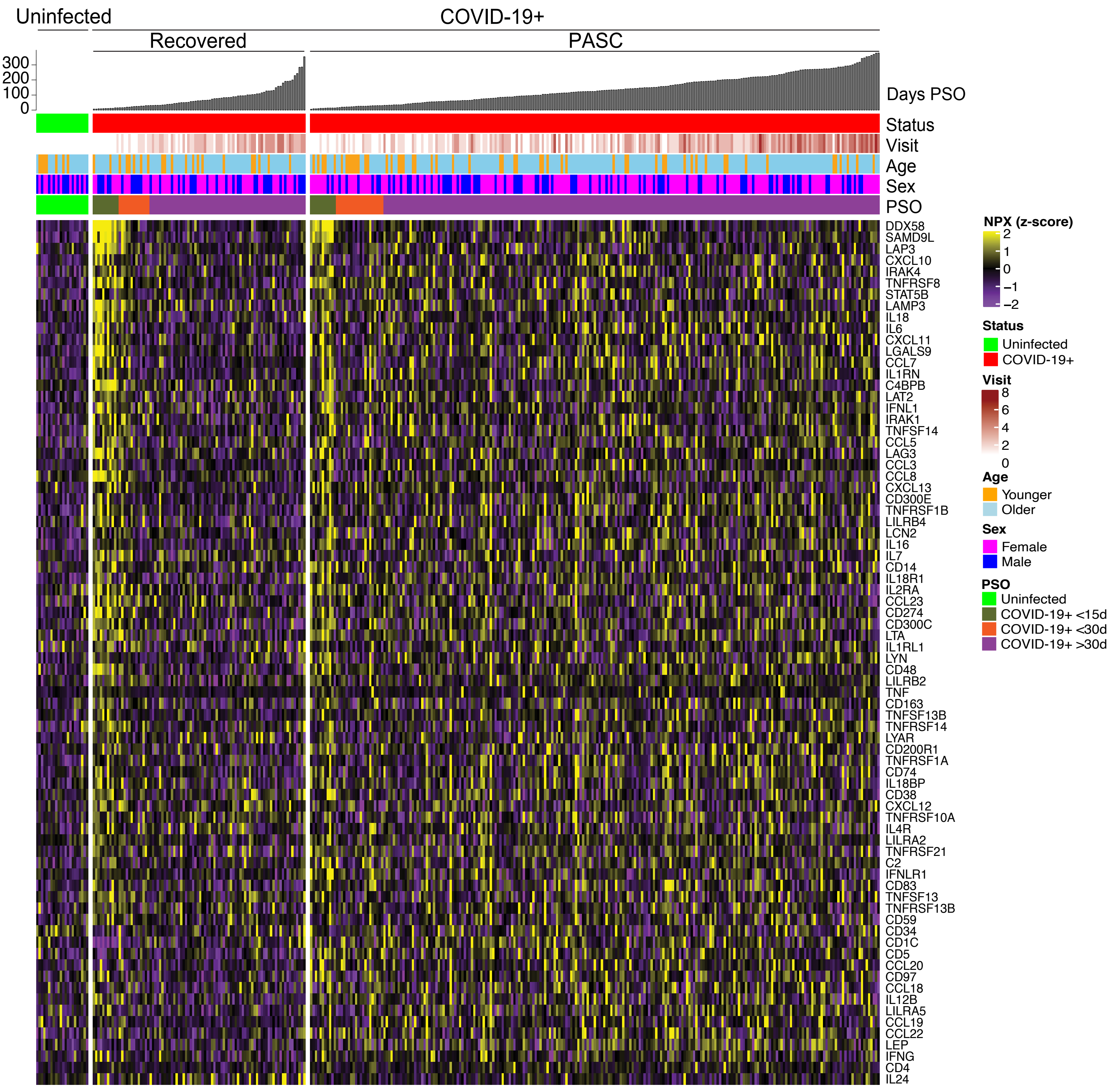

### Supplemental Figure S7

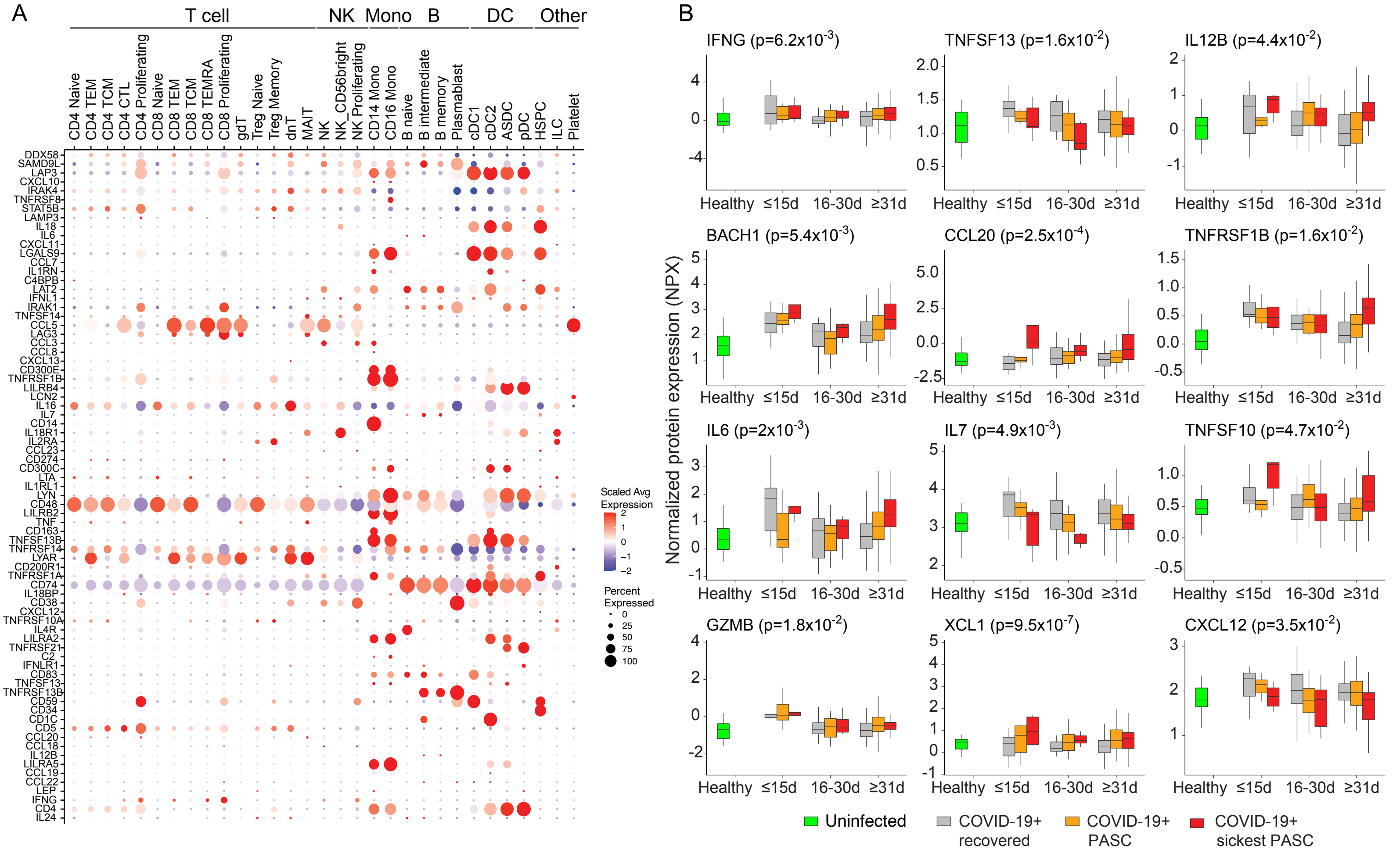
